# Exploring the antibiotic-producing capabilities of Canadian High Arctic bacteria

**DOI:** 10.64898/2026.10.11.757169

**Authors:** Eszter Farkas, Maja Lambeck, Grace Parish, Laurence Belanger, Daniella Serrador, Mina Nekouei, Kalinka Koteva, Francois Le Mauff, Patrick Łypaczewski, B Jesse Shapiro, Lyle Whyte, Gerard D Wright, Karine Auclair, Samantha Gruenheid, Dao Nguyen

## Abstract

Microbial communities in harsh environments, such as the Canadian High Arctic, remain a relatively unexplored source of antimicrobial natural products. In this study, we developed a pipeline for screening and evaluating bacterial isolates cultured from 23 Canadian High Arctic soil, sediment, and active-layer permafrost samples to discover antibacterial compounds. The pipeline incorporated an undergraduate studentsourcing program and produced a collection of over 350 bacterial isolates with preliminary antibacterial activity. Following dereplication, whole genome sequencing and secondary screening by co-culture assays, five isolates were prioritized to test crude extracts for antibacterial activity. Among bioactive crude extracts from these isolates, the *Pseudomonas* isolate with the most potent activity against *S. aureus* was analyzed. Using liquid chromatography bioactivity guided fractionation, we identified a potential bioactive compound known as koreenceine A, a compound previously only isolated from a *Pseudomonas* from temperate soil and involved in gene expression of other bacteria in their microbial community. Here we expanded the known antibiotic activity of koreenceine A and confirmed that a Δ*kecBCD* genetic knockout of the putative biosynthetic gene cluster significantly abrogated the antibacterial activity.

**Importance:** Environmental bacteria naturally produce antimicrobial compounds to compete and suppress other microbes around them. Polar bacteria are relatively understudied, especially from the Canadian High Arctic (CHA) region. In this study we report on the development of a pipeline which integrated a studentsourcing program to screen environmental samples from the CHA, followed by characterization of a collection of CHA bacterial isolates with antibacterial activity against human pathogens. This work contributes to a better understanding of the antibacterial-producing capabilities of Arctic bacteria.

## Background

Microbes from various environmental communities produce secondary metabolites to combat other bacteria in their environment (1). These compounds are commonly referred to as natural products (NPs), and many of them have historically been of interest to researchers for their antibacterial properties (2). The Canadian High Arctic (CHA) region is located above the Arctic circle and is known for its polar barrens, frigid temperatures, and high salt conditions (3). Studies on CHA ecological niches such as active-layer permafrost, permafrost, and hypersaline spring sediment have identified microbial communities that are unique to these locations (4–8). Understudied and harsh environments are of particular interest due to their potential to harbor microbes that produce compounds distinct from those found in temperate environments, and novel compounds have been isolated from several harsh environments, such as monorhamnolipids (Arctic ocean) (9), dithiolopyrrolones (Saharan desert) (10), and lanthipeptides (Georgian cave) (11). Isolation of bacteria from polar regions has resulted in the discovery of multiple unique NP with antibacterial activity (12–14).

The rise in antimicrobial resistance (AMR) and the dwindling of new compounds in the antibiotic development pipeline have converged to create a global AMR crisis that has put our ability to effectively treat infections at risk (15, 16). New antibiotics are required in the face of this crisis, and most antibiotics used clinically were isolated from soil bacteria during the “Golden Age” of discovery (17). Following the largely failed endeavours to discover new antibiotics from synthetic compounds, research efforts have returned to examining NPs from bacteria (18, 19). However, solation and screening of bacteria can be labour-intensive, especially with high volumes of isolates required for antibacterial drug discovery. To alleviate some of the labour-intensive process of culturing and screening soil bacterial isolates, researchers have turned to citizen science and student initiatives, such as the Tiny Earth program (20, 21) to crowdsource the search for novel antimicrobial NPs. Each year with Tiny Earth, thousands of students from over 500 academic institutions isolate bacteria from various sources and screen them for antibacterial activity, and to date have identified over 26,000 bacterial isolates that inhibit safe relatives of human pathogens (20, 21). After the initial identification of isolates with antibacterial activity, dereplication is essential to avoid the rediscovery of known antibiotics. To streamline the dereplication process, Cox et al developed the Antibiotic Resistance Platform (ARP), a simple and low-cost method which covers most major classes of antibiotics used in the clinic (22).

In this study, we developed a pipeline for the isolation, screening, and characterization of bacterial isolates from CHA soil, sediment, and active-layer permafrost samples. The pipeline integrated the crowdsourcing workflow developed by the Tiny Earth initiative for undergraduate students. Upon screening isolates from different CHA locations yielded, we created a collection of bacterial isolates from diverse genera that exhibited preliminary antibacterial activity. This collection was then processed through the screening pipeline to more extensively characterize their antibacterial activity.

## Methods

### Collection of soil, sediment and active-layer permafrost samples

Samples were collected during the summer from 23 locations (described in Table 1) near the McGill Arctic Research Station on Axel Heiberg Island, Nunavut, Canada, or near the Polar Continental Shelf Program facility, on Cornwallis Island, Nunavut, Canada (23) from habitats that include active layer permafrost, soil, and sediment. Samples were collected using an ethanol sterilized spatula, placed in sterile Whirl-Pak bags and kept at 5°C for transport and storage until testing. Some samples from active layer permafrost or Gypsum Hill Spring sediment were incubated *in situ* using a custom designed cryo-iPlate as described in Marcolefas et al (23). Briefly, 2% w/v gellan gum plugs within the plate were inoculated with 10µL of Arctic soil sample diluted in sterile water (∼1–10 microbial cells) and 0.03 μm-pore-size polycarbonate membranes (Whatman^TM^, GE Healthcare Life Sciences) were overlaid on the top and bottom of the plate. The plate was then incubated *in situ* for 7 days. The plugs were then transferred into Eppendorf tubes containing buffered glycerol solutions for storage until testing.

**Table 1.** Sampling location of Canadian High Arctic isolates.

| Habitat | Location | Habitat code | GPS coordinates | Number of antibacterial isolates from location | Habitat type |
| --- | --- | --- | --- | --- | --- |
| Ice wedge polygon permafrost at 2-inch depth | Axel Heiberg Island Nunavut | IWPP2 | N 74.43582<br>W 94.59688 | 20 | Active layer permafrost |
| Ice wedge polygon permafrost at 10-inch depth | Axel Heiberg Island Nunavut | IWPP10 | N 74.43582<br>W 094.59688 | 16 | Active layer permafrost |
| Arid active layer permafrost soil | Axel Heiberg Island Nunavut | AALPS | N 79.412872<br>W 90.740343 | 1 | Active layer permafrost |
| Active layer permafrost gravel soil | Axel Heiberg Island Nunavut | GSAL | N 79.412872<br>W 90.740343 | 15 | Active layer permafrost |
| Gypsum Hill hummock active layer permafrost soil | Axel Heiberg Island Nunavut | GHHS | Unknown | 106 | Active layer permafrost |
| Supplemented media active layer permafrost soil | Axel Heiberg Island Nunavut | iPAL | Unknown | 7 | Active layer permafrost |
| Polygon surface active layer permafrost soil | Axel Heiberg Island Nunavut | PSALP | N 79.412872<br>W 90.740343 | 1 | Active layer permafrost |
| Polygon wedge active layer permafrost soil | Axel Heiberg Island Nunavut | PWALP | N 79.412872<br>W 90.740343 | 2 | Active layer permafrost |
| Supplemented media active layer permafrost soil | Cornwallis Island | CIP | N 74.42995<br>W 95.0021 | 0 | Active layer permafrost |
| Assistance Bay Beach sediment (low tide) | Cornwallis Island, Nunavut | ALT | N 74.39046<br>W 94.17862 | 0 | Beach sediment |
| Assistance Bay Beach sediment (high tide) | Cornwallis Island, Nunavut | AHT | N 74.39050<br>W 94.17867 | 22 | Beach sediment |
| Dump Beach sediment | Cornwallis Island, Nunavut | DUBS | N 74.671699<br>W 94.922990 | 5 | Beach sediment |
| Dynamite Beach sediment | Cornwallis Island, Nunavut | DYBS | N 74.42143<br>W 95.04933 | 17 | Beach sediment |
| Tank Beach sediment | Cornwallis Island, Nunavut | TABS | N 74.41073<br>W 94.53328 | 6 | Beach sediment |
| Tupirvik Beach sediment | Cornwallis Island, Nunavut | TUBS | N 74.44610<br>W 95.02917 | 4 | Beach sediment |
| Arctic Ocean (salt water) sediment | Cornwallis Island, Nunavut | SW | Unknown | 10 | Sediment |
| Enriched gas bubble pond sediment | Cornwallis Island, Nunavut | GB | Unknown | 26 | Sediment |
| Freshwater lake sediment and biofilm | Cornwallis Island, Nunavut | FL | Unknown | 26 | Sediment |
| Gypsum Hill Spring outflow channel microbial mat | Axel Heiberg Island Nunavut | GHSM | N 79.403906<br>W 90.732321 | 3 | Sediment |
| Gypsum Hill Spring outflow channel sediment | Axel Heiberg Island Nunavut | GHSS | N 79.403906<br>W 90.732321 | 1 | Sediment |
| Supplemented media Gypsum Hill Spring cryo-plate plugs | Axel Heiberg Island Nunavut | iPGH | N 79.403906<br>W 90.732321 | 2 | Sediment |
| Arctic Willow (*Salix arctica*) rhizobium | Cornwallis Island, Nunavut | AWR | Unknown | 20 | Soil |
| Gravel soil | Cornwallis Island, Nunavut | GS | Unknown | 17 | Soil |
| Unknown | Unknown | Unknown | Unknown | 26 | Unknown |

### Isolation and screening of Arctic bacterial isolates using spread-patch assays

Arctic samples were processed as follows: briefly, 1 g of soil or 100µl of cryo-iPlate plug containing buffered solution was added to 9 mL of sterile water and serially diluted by 10-fold to 10^-7^. 100 µL of each sample dilution was plated on Tryptic Soy Agar (TSA) (BBL BD) plates containing 100 µg/mL of amphotericin B and incubated at room temperature (23°C). After 1 week, morphologically distinct colonies were selected based on colour, size, and shape. These Arctic isolates were screened for antibacterial activity against non-pathogenic relatives of human pathogens (Table S1), namely *Enterococcus raffinosus, Bacillus subtilis, Staphylococcus epidermidis, Escherichia coli, Acinetobacter baylyi, Pseudomonas putida,* and *Enterobacter aerogenes* using the spread-patch assay as follows: for each pathogen-relative tester species, cultures were grown in 5 mL Tryptic Soy Broth (TSB) (Bacto BD) for 18 h at 37° C without shaking, and 150 µL liquid culture (∼10^8^ CFU) was evenly spread on TSA plates to form a lawn, with each plate containing one pathogen-relative tester species. Arctic isolates were then patched from colonies onto these plates, with approximately 24 Arctic isolates per plate. After one week of incubation at room temperature, the zone of inhibition (ZOI) surrounding the patch of Arctic isolates was observed, indicating inhibition of the tester species by the Arctic isolate. Arctic isolates that produced a ZOI of at least 1 mm against at least one pathogen-relative tester species were considered to have antibacterial activity, selected, and stored as glycerol freezer stocks for further characterization.

### 16S rRNA gene sequencing of Arctic isolates

Colony PCR to amplify the 16S rRNA gene was performed on all Arctic isolates that inhibited at least one pathogen-relative. The PCR reaction used primers (27F forward primer: 5′-AGAGTTACCTTGTTACGACTT-3′ and 1492R reverse primer: 5′-GGTTACCTTGTTACGACTT-3′) with the following cycling program: (1) 94°C for 10 min, (2) 30 cycles of 94°C for 30 s, 58°C for 30 s, 72°C for 1 min, (3) 72°C for 10 min. PCR products were sequenced using Sanger sequencing technology at the Génome Québec Innovation Centre. The sequences were assessed for quality using 4Peaks software (24). The species of the isolates was identified using nucleotide BLAST, where isolates with less than 97% match to known species in the 16S ribosomal RNA sequences (Bacteria and Archaea) database were considered to be new species (Table S2).

### Co-culture stamping assay

Arctic isolates with antibacterial activity against at least one pathogen-relative tester species were selected for subsequent testing against ARP strains (for dereplication) and human pathogenic strains. Bacteria were grown in 5 mL of LB Miller media (Difco) for 18 h at 37°C with shaking at 250 RPM. Cultures were diluted in sterile phosphate buffered saline (PBS) (Wisent) to an OD_600_ of 0.1, and 300 µL of the diluted culture was spread on TSA agar (Wisent) rectangular plates immediately prior to stamping of Arctic isolates onto the sample plate. Arctic isolates were first grown in 96-well microtiter plates in 200 µL of TSB media for 18 h at 23°C with shaking at 250 RPM. Approximately 2 µL of Arctic isolates cultures were stamped from the 96-well plates onto TSA rectangular plates using a sterile 96-tinged stamper. ZOI were observed surrounding the patch of Arctic isolates after 1 week of incubation at 23°C, indicating inhibition of the tester strain by the Arctic isolate. ZOI were designated as 3 (large), 2 (medium), 1 (small), or 0 (none) based on visual assessment (Figure S1).

### ARP dereplication

A selected ARP strain panel (Table S3) representing most antibiotic classes used in the clinic was used for dereplication using the co-culture stamping assay (22). All ARP strains are in the hyperpermeable, efflux-deficient *E. coli* BW25113 Δ*bamB* Δ*tolC* (25) background except for the *E. coli* pGDP1 MCR-1 strain with polymyxin resistance in the wild-type *E. coli* BW25113 background (22). An Arctic isolate was considered to “pass” dereplication if it inhibited all of the ARP strains with a ZOI of at least 1 mm. Isolates that did not inhibit the MCR-1-expressing polymyxin resistant strain were only considered to have passed dereplication if they also failed to inhibit the parental wild-type *E. coli* BW25113 strain.

### Whole genome sequencing (WGS) and analysis

Full sequencing methods can be found in the supplemental methods. Briefly, genomic DNA was purified using the DNeasy Blood & Tissue Kit (Qiagen) according to the manufacturer’s protocol. Hybrid WGS (Illumina and Oxford Nanopore Technologies (ONT)) or long-read sequencing (Nanopore Rapid Barcoding 96 v14, ONT, with v4.3@sup models) was performed for 2018 isolates and MNAAK/2019/2021 isolates respectively. Assembly was performed using tools available in the open source, web-based platform Galaxy (https://usegalaxy.org/) with *de novo* assembly using Unicycler (v 0.5.0+galaxy1) (26) or provided by Plasmidsaurus.

Whole genome sequences of Arctic isolates were compared for average nucleotide identity (ANI) using FastANI (27). Isolates with ANI of 99.99% were considered identical strains and only one of the isolates was retained for further characterization. All isolates with ANI < 99.99% were considered different strains and continued in the pipeline (Table S4). Taxonomic classification was done using the Microbial Genomes Atlas (MiGA) (28).

### Genome analysis for biosynthetic gene clusters (BGCs)

WGS assemblies of Arctic isolates were then analyzed with the antibiotics and small molecules analysis shell (antiSMASH, version 8.0.4) open-source software via web access (29). Analyses were done using both the “relaxed” setting (which detects homology with complete and partial well-defined clusters) and the “loose” setting (which also detects homology with poorly-defined clusters), and included the features of KnownClusterBlast, MIBiG cluster comparison, Cluster Pfam analysis, TFBS analysis, ClusterBlast, ActiveSiteFinder, SubClusterBlast, and RREFinder.

### Preparation of crude extracts from Arctic isolate cultures

Cultures of Arctic isolates were grown in 15 mL of TSB medium (125 mL baffled flasks) for 24 h at 23 °C. Cultures were centrifuged at 7200 x g for 15 min. The culture supernatant was then collected and extracted with 1:1 ethyl acetate (EtOAc) (Fisher Scientific) three times. With each cycle of extraction, the aqueous and organic phases were allowed to separate and collected separately. The organic layer was dried using a rotary evaporator (R0215 Rotavapor, Buchi) to generate crude extracts. Crude extracts were named as CE plus the isolate ID from which it was generated (CE_isolate ID).

### Well diffusion assay for antibacterial activity of crude extracts

Human pathogen tester strains (listed in Table S5) were grown in 5 mL of TSB media for 18 h at 37°C. Cultures were diluted in sterile PBS to an OD_600_ of 0.1, and 200 µL of the bacterial suspension was evenly spread on TSA agar plates. 6 mm wells were then punched into the agar plates. Crude extracts were resuspended in 100% HPLC-grade methanol (MeOH) (Sigma Aldrich) to a concentration of 166.67 mg/mL (for MNAAK) or 333.33 mg/mL (all others). 30 µL of crude extract (5 mg CE_MNAAK, 10 mg for all others) were pipetted into each well of the agar plates. Plates were left to dry for 30 min then incubated for 24 h at 37°C. ZOI were measured as the total diameter of the ZOI minus the diameter of the agar.

### Bioactivity guided fractionation of CE_MNAAK

The Arctic isolate MNAAK was grown under optimized conditions in 500 mL of R2A media (Teknova) in a 2 L flask at 15°C for 24 h with shaking at 250 RPM. Following the established extraction protocol described above, CE_MNAAK was first fractionated by Preparative-Thin Layer Chromatography (Prep-TLC) using Supelco glass-backed silica gel plates (Sigma Aldrich) with a solvent system consisting of 47.5% dichloromethane (DCM): 47.5% EtOAc: 5% MeOH + 0.1% formic acid. Fractions were labelled starting with the most polar fraction on the bottom (fraction 1). Fractions were tested in a bioluminescent *E. coli* assay as described in Farkas et al. 2025 (30).

To further fractionate fraction 1, semi-preparative high-performance liquid chromatography (HPLC) was conducted using an Agilent 1100 series system (Agilent Technologies) with a diode array measuring absorbance at 214 nm and a fraction collector using a Luna C18 column (Phenomenex 250 mm × 10 mm × 5 μM particle size). To do so, 50 μL of sample (∼100 mg/mL) was injected at a 4.5 mL/min flow rate. Separation was performed using a linear gradient of 1-99% acetonitrile (ACN) + 0.1 % formic acid in water over 50 min. The solvent of each fraction was evaporated under reduced pressure before lyophilisation to remove residual water.

Each fraction was tested in a microdilution minimal inhibitory concentration (MIC) assay. Briefly, lyophilized compounds from each fraction were resuspended in cation-adjusted Mueller-Hinton broth (CAMHB, BD) media with 2.5% MeOH to a concentration of 2048 µg/mL. Next, 100 µL was added to the empty wells in the last column of a 96-well plate and serially diluted 2-fold in 50 µL of CAMHB media, leaving the first column empty as negative control. 50 µL *of E. coli Δbam ΔtolC* diluted to an OD_600_ of 0.001 in CAMHB media was added to each well, resulting in a final concentration of 1.25% MeOH and 1024 to 1 µg/mL of crude extract in the assay. The MIC was determined by visual inspection after 20 hrs of static incubation at 37 °C with a 15 min shaking incubation (200 RPM at 37°C) at hour 18 (31).

Liquid chromatography coupled to low resolution mass spectrometry (LC-LRMS) analysis (scan range 100–2000 *m/z,* diode array measurement at 214 nm) of fraction 1 from the CE_MNAAK crude extract was performed using the Agilent 1100 series HPLC system and Agilent 6120 single quadrupole mass spectrometer (Agilent technologies) with a heated electrospray ionisation (ESI) source (in positive ESI mode) using a Luna C18 column (Phenomenex 250 mm 4.6 mm × 5 μM particle size). To do so, 10 μL of sample at a concentration of 10 mg/mL was injected at a flow rate of 1 mL/min under the same solvent settings as for semi-preparative HPLC.

In a second stage, CE_MNAAK was fractionated by flash chromatography using the CombiFlash Rf* Lumen (Teledyne ISCO) using a RediSep Gold 15.5 g HP C18 column (Teledyne ISCO). The crude extract (∼120 mg) was loaded onto the column, and separation was performed at a flow rate of 30 ml/min using a solvent system of H_2_O and ACN. The methods consisted of a hold of 10% ACN for 3 min, then a linear gradient of 1–100 % solvent ACN over 14 min, followed by a hold of 5 min at 100 % solvent ACN. Fractions were collected and dried overnight by evaporation using the SP Genevac (Canadawide Scientific).

Each fraction was tested in an overnight activity assay against *E. coli* Δ*bamB* Δ*tolC*, *S. aureus* ATCC 29213, and *A. baumannii* ATCC 17978. Overnight cultures of tester pathogens were diluted to an OD_600_ of 0.0005 and 98 µL was added to each well of a 96-well plate. Dried fractions were resuspended in 200 µL DMSO and 2µL was added to the 98µL of pathogen. The plate was then incubated for 18 h at 37°C. The OD_600_ of each well in the assay plate was then read using the Synergy H1 microplate reader (Biotek).

Liquid Chromatography – High Resolution Mass Spectrometry (LC-HRMS) analyses of the active fraction (scan range 100–3000 *m/z,* diode array measurement at 214 nm) were performed using the 6550 iFunnel Q-TOF LC-MS (Agilent technologies), equipped with 1290 Infinity II UPLC (Agilent Technologies) in positive ESI mode using a Zorbax Eclipse XDB C8 column (Agilent 100mm x 2.1 mm x 3.5 µm particle size. To do so, 10 μL of sample injected at a flow rate of 0.4 mL/min with a linear gradient of 5% to 95% ACN over 10 min.

### Construction of the *P. prosekii* MNAAK Δ*kecBCD* mutant

The *ΔkecBCD* unmarked in-frame deletion construct was generated by commercially synthesizing a chimeric DNA fragment (GeneArt™ Gene Synthesis. Thermo Fisher Scientific) consisting of the 500 bp region upstream of the *kecB* gene and the first 9 bp of the *kecB* open reading frame (ORF) fused with the last 9 bp of the *kecD* ORF and the 500 bp region downstream of the *kecD* gene from the *P. prosekii* MNAAK. The fragment was then inserted into the pDONRpEX18-Gm suicide vector (32) at the *attB* site using Gateway cloning methods (33).

The *P. prosekii* MNAAK Δ*kecBCD* knockout mutant was generated by a two-step allelic exchange approach (32). Briefly, the pEX18-Gm-Δ*kecBCD* plasmid was first transformed into the *E. coli* SM10 conjugative strain via electroporation, and transformants were selected with 10 µg/mL gentamicin. Biparental mating was performed according to Hmelo et al (32) using the *E. coli* SM10 pEX18-Gm-Δ*kecBCD* strain (growth in 50 mL of LB Miller broth supplemented with 10 µg/mL gentamicin for 18 h at 37°C, 200 RPM) and the *P. prosekii* MNAAK strain (growth in 15 mL of TSB broth for 18 h at 23°C, 200 RPM). Cells from both cultures were then pelleted by centrifugation, mixed, spotted on a TSA plate, and incubated for 24 h at 23°C. Cells were then resuspended in sterile PBS and plated on TSA plates containing 50 µg/mL gentamicin and 10 µg/mL chloramphenicol for selection of merodiploids. Transformants were then re-streaked on TSA plates containing 15% sucrose for counterselection of gentamicin susceptible clones. The *P. prosekii* MNAAK Δ*kecBCD* mutant was confirmed by whole genome sequencing.

## Results

### Sampling, isolation, and initial screening of Arctic isolates for antibacterial activity

Building on an initial pilot study on Arctic bacteria reported by Marcofelas et al (23), we integrated the workflow of undergraduate students at McGill participating in the Tiny Earth program (21) to create a Canadian High Arctic (CHA) bacterial isolate collection. The pipeline was optimized to broaden the range of environmental samples, expand the panel of human pathogen tester strains, and include characterization and identification of bioactive compounds. The schematic representation of our study’s pipeline is outlined in Figure 1. Following sample collection, bacteria were isolated from the soil/sediment/ active-layer permafrost samples by culture, screened for initial antibacterial activity against pathogen-relative tester strains, and dereplicated. Unique Arctic isolates were prioritized based on antibacterial activity against human pathogens, with further analysis of the bacterial extracts to identify bioactive compounds.

**Figure 1.**
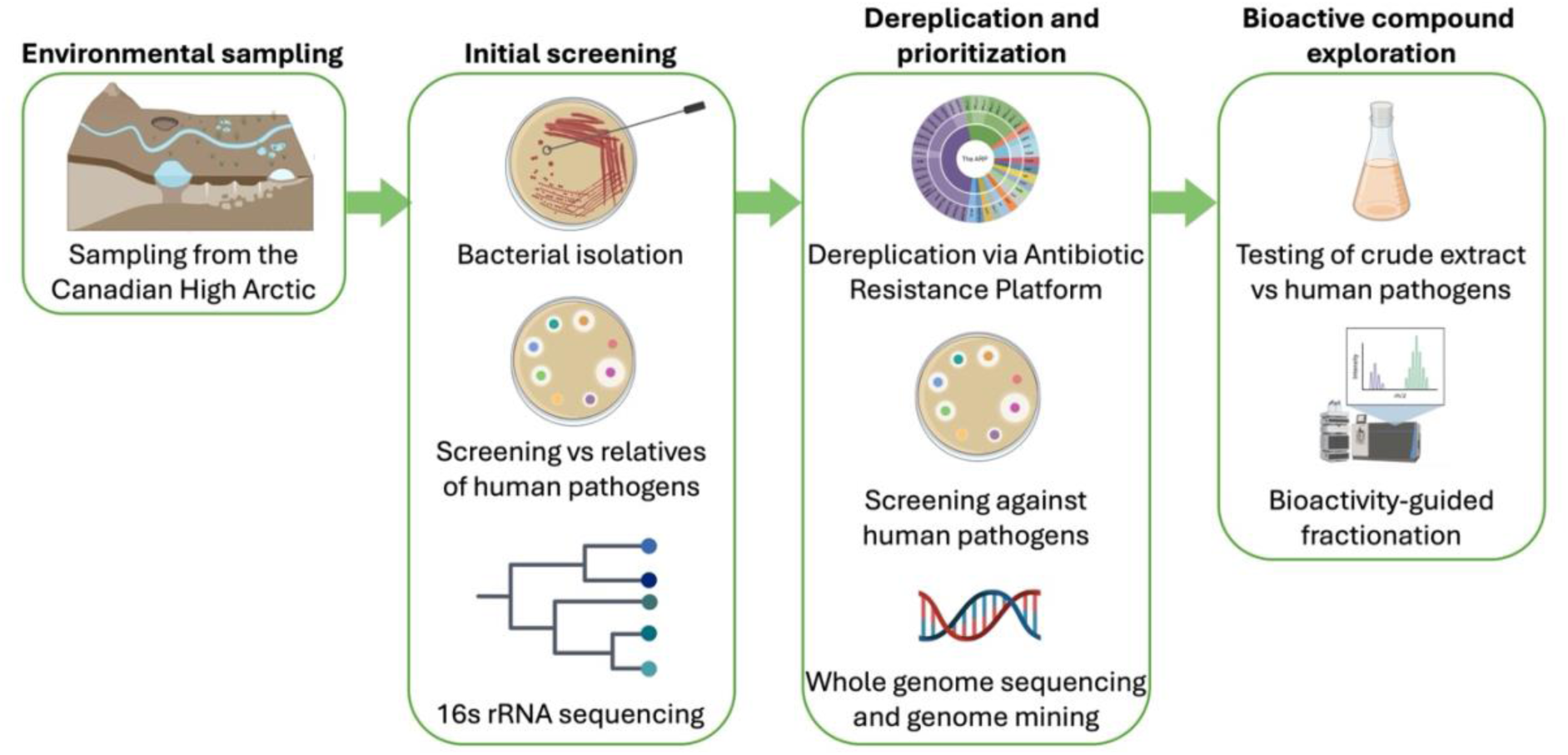
Schematic representation of the antibiotic discovery pipeline for this study. Samples from 23 locations from active layer permafrost, soil, beach sediment, and sediment habitats were collected. Initial screening steps were performed by students in an undergraduate microbiology teaching lab as part of the Tiny Earth initiative. Isolates that passed the initial screening step were put through a dereplication step, confirmed for activity against human pathogens, and WGS to filter out duplicate strains, resulting in 11 candidate strains, of which five were prioritized. Ethyl acetate crude extracts from the five prioritized bacterial isolates were tested for activity against *S. aureus* strains and one crude extract was prioritized for bioactivity guided fractionation. Created in BioRender. Farkas, E. (2026) https://BioRender.com/iyglqu9

The McGill Arctic Research Station (MARS) is located in the CHA at Expedition Fjord on Axel Heiberg Island (Figure 2A), an area that includes perennial springs, polygonal terrain, and a polar desert climate. The Polar Continental Shelf Program (PCSP) facility is located on Cornwallis Island in the CHA (Figure 2A) and experiences a polar desert climate (34). Samples were collected from 23 locations (listed in Table 1) belonging to sediment, soil, and active-layer permafrost habitats near the MARS and PCSP. Sediment samples included lake, ocean, spring, and beach sediments. Active-layer permafrost samples of gravel soil, hummock soil, polygonal terrain, and ice wedge polygon soil. Soil samples were from the rhizosphere of the Arctic Willow (*Salix arctica*) and gravel soil.

**Figure 2.**
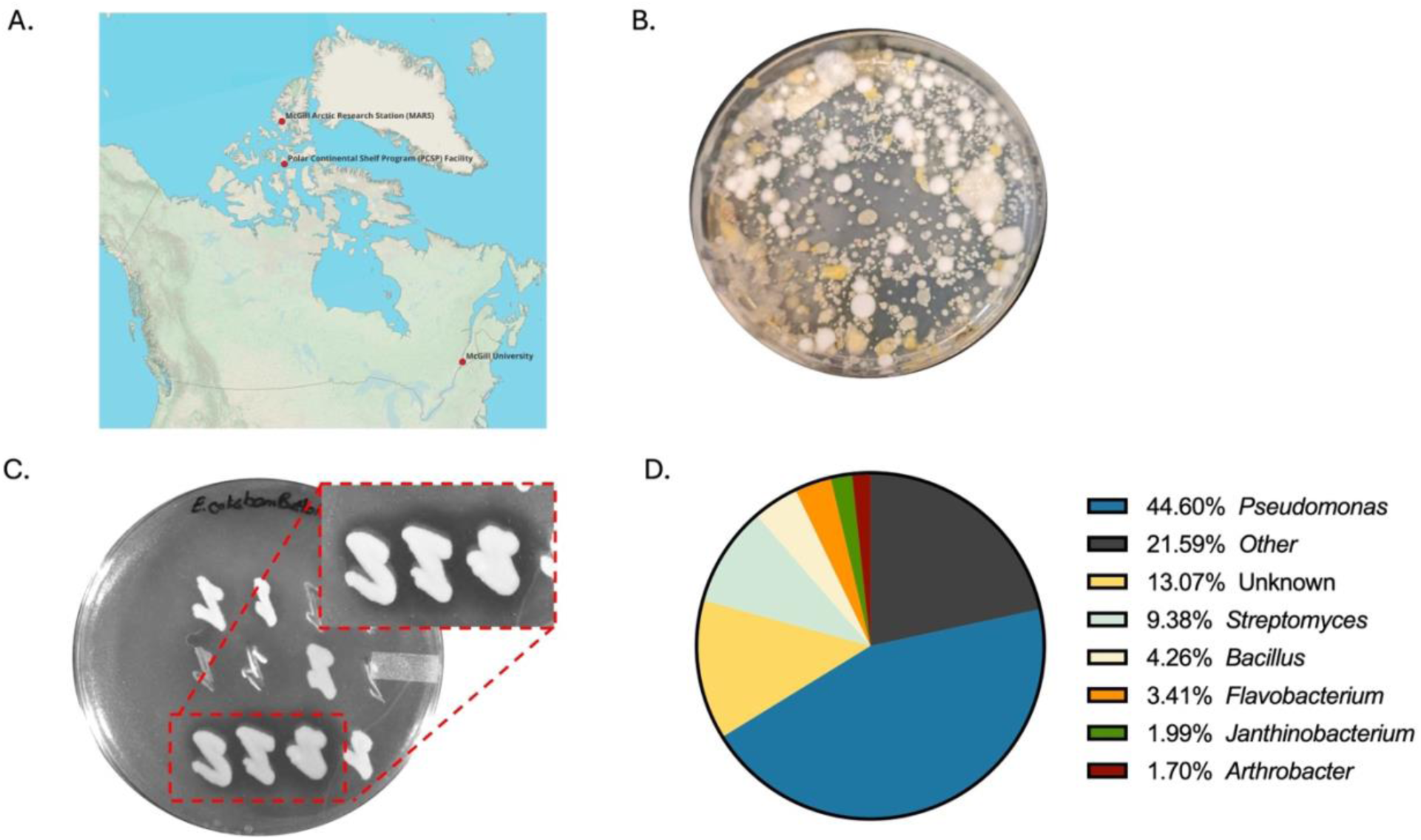
Isolation of bacterial isolates from CHA soil, sediment and active-layer permafrost samples and screening against safe relatives of ESKAPE pathogens. **A.** Location of McGill Arctic Research Station on Axel Heiberg Island, Polar Continental Shelf Program facility on Cornwallis Island, and McGill University. Map created using QGIS open-source software (35). **B.** Representative image of soil suspension cultured on TSA plates containing amphotericin B for isolation of Arctic bacterial isolates. **C.** Representative image of spread-patch assays to test Arctic isolates’ activity against pathogen-relative tester species. A lawn of each tester strain was spread on a TSA plate, on top of which individual Arctic isolates were patched. The ZOI was assessed after incubation for 1 week at room temperature. Inset highlights isolates displaying a ZOI. Image contrast enhanced for clarity. **D.** Taxonomic identification by 16S rRNA gene sequencing of Arctic isolates with activity against pathogen-relative tester species.

Over 20,000 bacterial colonies were isolated by culture-based methods from Arctic samples (Figure 2B) by approximately 140 students. For a simple phenotypic readout to prioritize these isolates, they were tested for antibacterial activity against human pathogen safe relatives (Figure 2C) using the spread-patch assay. A total of 352 isolates inhibited at least one pathogen-relative tester species (*Bacillus subtilis, Staphylococcus epidermidis, Acinetobacter baylyi, Pseudomonas putida, Enterobacter aerogenes, Escherichia coli,* Table S2) based on a visible ZOI on TSA plates. Isolates with antibacterial activity were identified from 21 of the 23 locations sampled (Table 1). To characterize the taxonomic diversity of the CHA isolate collection, we used 16S rRNA gene sequencing to identify the genera of isolates. The genera of 316 of the 352 isolates were successfully identified, as shown in Figure 2D & Table S2, with the largest group (44.60%) belonging to the *Pseudomonas* genus, followed by *Streptomyces* (9.38%). *Bacillus* (4.26%), *Flavobacterium* (3.41%), *Janthinobacterium* (1.99%), and *Arthrobacter* (1.70%). The remaining 21.59% of isolates belong to other genera. While *Pseudomonas* isolates were isolated from most locations, *Streptomyces* isolates were only isolated from three locations, with the majority from Gypsum Hill hummock active-layer permafrost soil (Figure S2).

### Dereplication using the Antibiotic Resistance Platform

To avoid the rediscovery of compounds chemically similar to known antibiotics, Arctic isolates that inhibited at least one pathogen-relative tester species were put through a dereplication process using a subset of the Antibiotic Resistance Platform (ARP) (22). We used an ARP panel that consisted of 17 *E. coli* strains, each expressing a plasmid-encoded resistance gene to one of 16 distinct antibiotic classes (Table S3). Arctic isolates were screened against the ARP strains in co-culture stamping assays. An isolate was considered to “pass” dereplication and to produce bioactive compound(s) likely distinct from known antibiotics if it inhibited all the ARP strains. 138 isolates did not inhibit the *E. coli ΔbamB ΔtolC* ARP parental strain, thus could not be assessed by the ARP, and 47 isolates did not grow following initial isolation. From the 167 Arctic isolates successfully assessed with the ARP panel, 18 passed the dereplication process.

### WGS and biosynthetic capabilities of Arctic isolates

To avoid duplicate strains in subsequent steps of our pipeline, WGS was performed on the 18 Arctic isolates that passed dereplication. The average nucleotide identity (ANI) between all isolate pairs was determined using FastANI (27). Using a threshold of ANI >99.99% to define and exclude identical strains, a total of 11 unique isolates were selected for further characterization (Table S4). We then employed the Microbial Genomes Atlas (28) for taxonomic classification and identified the closest-matching known species to each Arctic isolate. Six isolates were identified as *Pseudomonas* species (closest relatives: *P. mandelii*, *P. prosekii*, *P. wadenswilerensis*, *P. parakoreensis*, *P. rossensis*, *P. baetica*) as *Serratia* species (closest relatives: *S. marcescens* and *S. ureilytica),* and two as *Bacillus* species (closest relatives: *B. pumilus* and *B. subtilis*), as shown in Table 2.

**Table 2.**
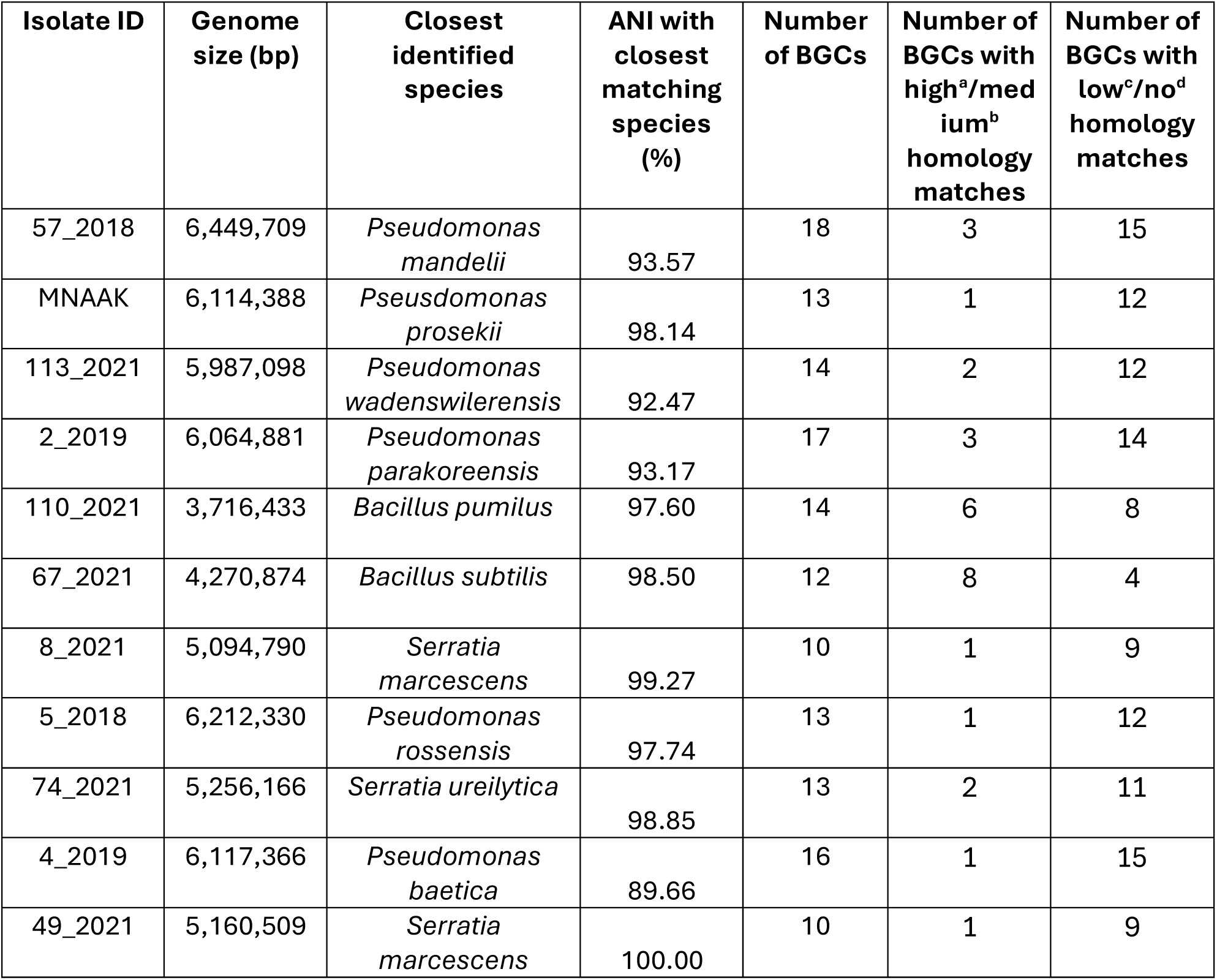
Genomic information on candidate Arctic isolates. ANI: Average nucleotide identity. ANI with closest matching species was performed using MiGA. BGCs identified using antiSMASH v8.0.4 software under “relaxed” setting. ^a^High homology match to known clusters: cluster similarity of larger or equal to 75%. ^b^Medium homology match to known clusters: cluster similarity between 75% and 50%. ^c^Low homology match to known clusters: cluster similarity between 50 and 15%. ^d^No homology match to known clusters: cluster similarity < 15%.

| Isolate ID | Genome size (bp) | Closest identified species | ANI with closest matching species (%) | Number of BGCs | Number of BGCs with high<sup>a</sup>/medium<sup>b</sup> homology matches | Number of BGCs with low<sup>c</sup>/no<sup>d</sup> homology matches |
| --- | --- | --- | --- | --- | --- | --- |
| 57_2018 | 6,449,709 | *Pseudomonas mandelii* | 93.57 | 18 | 3 | 15 |
| MNAAK | 6,114,388 | *Pseudomonas prosekii* | 98.14 | 13 | 1 | 12 |
| 113_2021 | 5,987,098 | *Pseudomonas wadenswilerensis* | 92.47 | 14 | 2 | 12 |
| 2_2019 | 6,064,881 | *Pseudomonas parakoreensis* | 93.17 | 17 | 3 | 14 |
| 110_2021 | 3,716,433 | *Bacillus pumilus* | 97.60 | 14 | 6 | 8 |
| 67_2021 | 4,270,874 | *Bacillus subtilis* | 98.50 | 12 | 8 | 4 |
| 8_2021 | 5,094,790 | *Serratia marcescens* | 99.27 | 10 | 1 | 9 |
| 5_2018 | 6,212,330 | *Pseudomonas rossensis* | 97.74 | 13 | 1 | 12 |
| 74_2021 | 5,256,166 | *Serratia ureilytica* | 98.85 | 13 | 2 | 11 |
| 4_2019 | 6,117,366 | *Pseudomonas baetica* | 89.66 | 16 | 1 | 15 |
| 49_2021 | 5,160,509 | *Serratia marcescens* | 100.00 | 10 | 1 | 9 |

Next, we examined the genomes for biosynthetic gene clusters (BGCs) associated with antibiotic production using antiSMASH (29), a leading bioinformatic tool which allows for identification, annotation, and prediction of BGCs based on an experimentally validated database and manually curated rules of secondary metabolite BGCs. From the 11 unique Arctic isolates selected, a total of 150 BGCs were identified, with an average of 14 BGCs per genome (range 10 to 18), with *Pseudomonas* isolates harbouring the highest number of predicted BGCs (Table 1). The BGCs classes (as shown in Figure S3) included a predominance of hybrid-type BGCs (25 of 150), followed by non-ribosomal peptide synthetase (NRPS, 20 of 150) and ribosomally synthesized and post-translationally modified peptide (RiPP)-like clusters (20 of 150). Of the 150 BGCs, 121 had low (15 to 50%) or no (<15%) homology to known BGCs based on sequence similarity), suggesting that these predicted BGCs might produce potentially novel compounds (Table 2).

### Screening for activity against human pathogens

To confirm the antibacterial activity against human pathogens (Table S5), the 11 Arctic isolates (Table S6) were tested against at least two different strains of four human pathogenic species (*S. aureus*, *A. baumannii*, *K. pneumoniae*, *P. aeruginosa*) selected because they are among the highest causes of healthcare-associated infections and the WHO Priority Pathogens List (36, 37). The antibacterial activity was measured in a co-culture assay by the ZOI for each Arctic isolate against the tester pathogen (Figure 3B), and *E. coli*Δ*bamB* Δ*tolC* was used a positive control. As shown in Figure 3A, 10 of the 11 isolates displayed activity against at least one human pathogenic strain, and 5 isolates displayed activity against multiple pathogenic species. Isolates 4_2019, 74_2019, 8_2021, 67_2021, and 110_2021 had antibacterial activity limited to *S. aureus*, while 57_2018 displayed the broadest spectrum of activity, with inhibition of all four *S. aureus* strains, both *A. baumannii* strains, and one of two *K. pneumoniae* strains tested. Isolates MNAAK, 113_2021, 2_2019, and 8_2021 displayed varying levels of antibacterial activity against all four *S. aureus* strains as well as *A. baumannii* ATCC 19606. Isolate 49_2021 displayed the most modest activity, with activity only against the *E. coli* Δ*bamB* Δ*tolC* control strain. None of the isolates inhibited *K. pneumoniae* ATCC 700603 or either of the *P. aeruginosa* strains.

**Figure 3.**
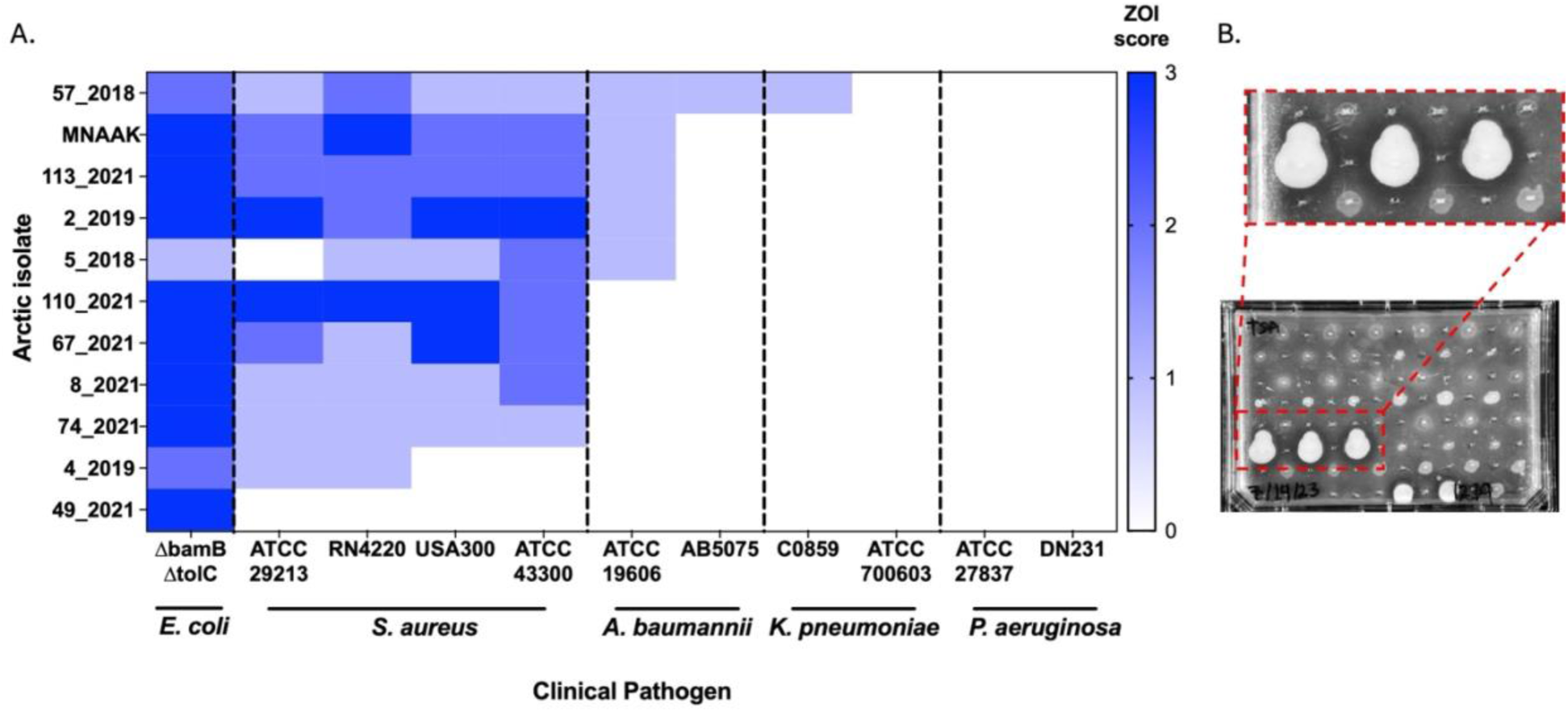
Antibacterial activity of Arctic isolates against human pathogens in co-culture stamping assay. **A.** Antibacterial activity of 11 unique Arctic isolates prioritized based on ARP dereplication and WGS. Antibacterial activity was measured in co-culture stamping assay by ZOI, ranked as 3 (large), 2 (medium), and 1 (small), or 0 (none) (scoring scale shown in Figure S1). Results are shown as the mode of N≥6 replicates. **B.** Representative image of stamping assay with inset highlighting the observable ZOI. Image contrast enhanced for clarity.

### Testing of crude extracts

Following co-culture testing, we prioritized five Arctic isolates that displayed antibacterial activity against multiple pathogenic species to characterize the bioactivity of secreted compound(s). As a first step, crude extracts were generated from culture supernatants using ethyl acetate to extract a wide range of polar secreted compounds (38). The crude extracts were then tested for antibacterial activity against four *S. aureus* strains and *E. coli ΔbamB ΔtolC* (as control) in a well-diffusion assay (Figure 4ACB). All extracts except for CE_5_2018 exhibited activity against the *E. coli ΔbamB ΔtolC* tester strain. CE_2_2019 and CE_113_2021 exhibited antibacterial activity against all four *S. aureus* strains tested at a concentration of 10 mg/well (Figure 4A). CE_57_2018 displayed strong activity against the *E. coli ΔbamB ΔtolC* strain but produced the smallest ZOI against all *S. aureus* strains (Figure 4A). Although CE_MNAAK did not inhibit methicillin-resistant *S. aureus* (MRSA) strain ATTC 43300, it showed greatest potency against the other *S. aureus* strains tested, leading us to prioritize this extract for further characterization.

**Figure 4.**
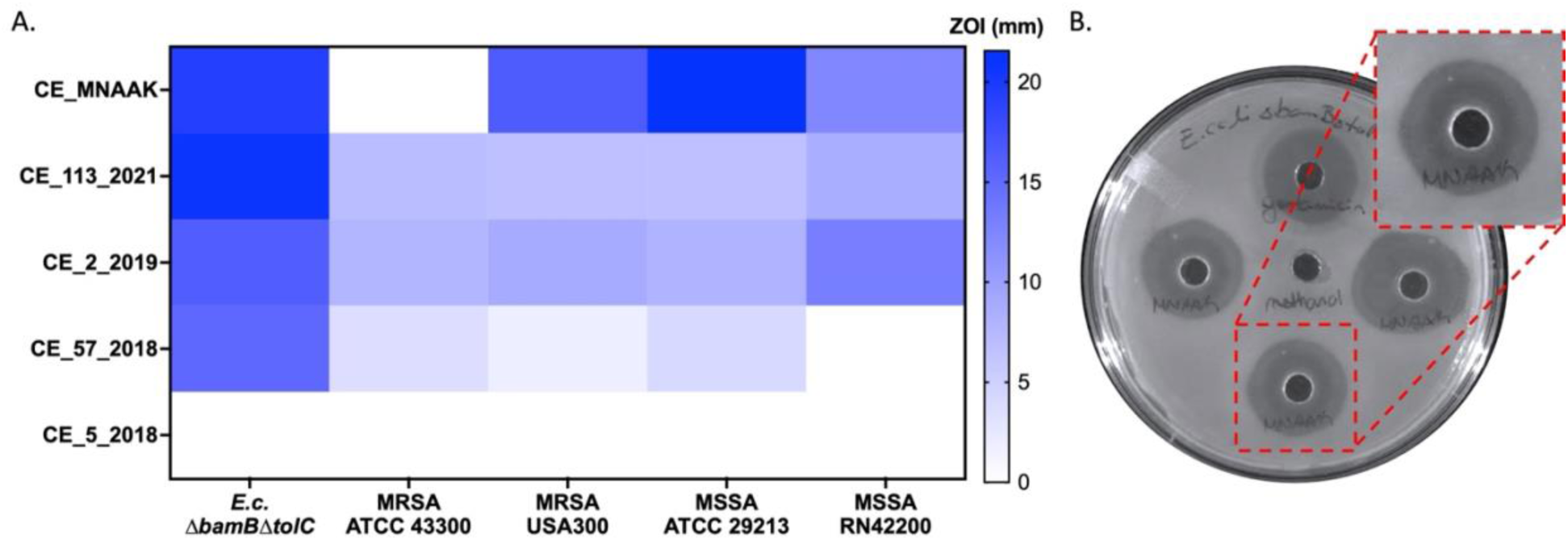
Antibacterial activity of prioritized crude extracts in well-diffusion assay. Ethyl acetate crude extracts from bacterial supernatant were tested at 5 mg/ well (CE_MNAAK) or 10 mg/ well (CE_5_2018, CE_57_2018, CE_2_2019, CE_113_2021) against the *E. coli ΔbamB ΔtolC* and two methicillin-susceptible (MSSA) and two methicillin resistant (MRSA) *S. aureus* strains in well diffusion assays. **A.** Antibacterial activity of crude extracts generated from 5 prioritized Arctic isolates against human pathogens. Results are shown as the mode of N≥6 replicates pooled from ≥2 independent experiments. **B**. Representative image of well-diffusion assay with observable ZOI. Image contrast enhanced for clarity.

### Bioactivity-guided fractionation of CE_MNAAK

In order to determine the compound(s) responsible for the antibacterial activity of CE_MNAAK, the crude extract was fractionated in an iterative manner using bioactivity-guided fractionation. The crude extract was first fractionated into 14 fractions using Preparative-Thin Layer Chromatography (Prep-TLC). Each fraction was tested for inhibitory activity using a bioluminescent *E. coli* strain, and several fractions displayed activity reported in Farkas et al. 2025 (30). Fraction 1, which contained the most polar compounds, was further fractionated into 17 fractions using HPLC with an acetonitrile (ACN) and water (H_2_O) gradient (Figure 5). These fractions (1.1 to 1.17) were then tested for activity against the *E. coli ΔbamB ΔtolC* strain in a MIC assay, and Fraction 1.7 displayed the greatest activity with a MIC of 16 mg/mL (Figure 5). Based on this fraction’s Low-Resolution Mass Spectrometry (LRMS) total ion chromatogram, we identified a peak containing a putative compound with a mass-to-charge (m/z) ratio of 208.3 (Figure S4).

**Figure 5.**
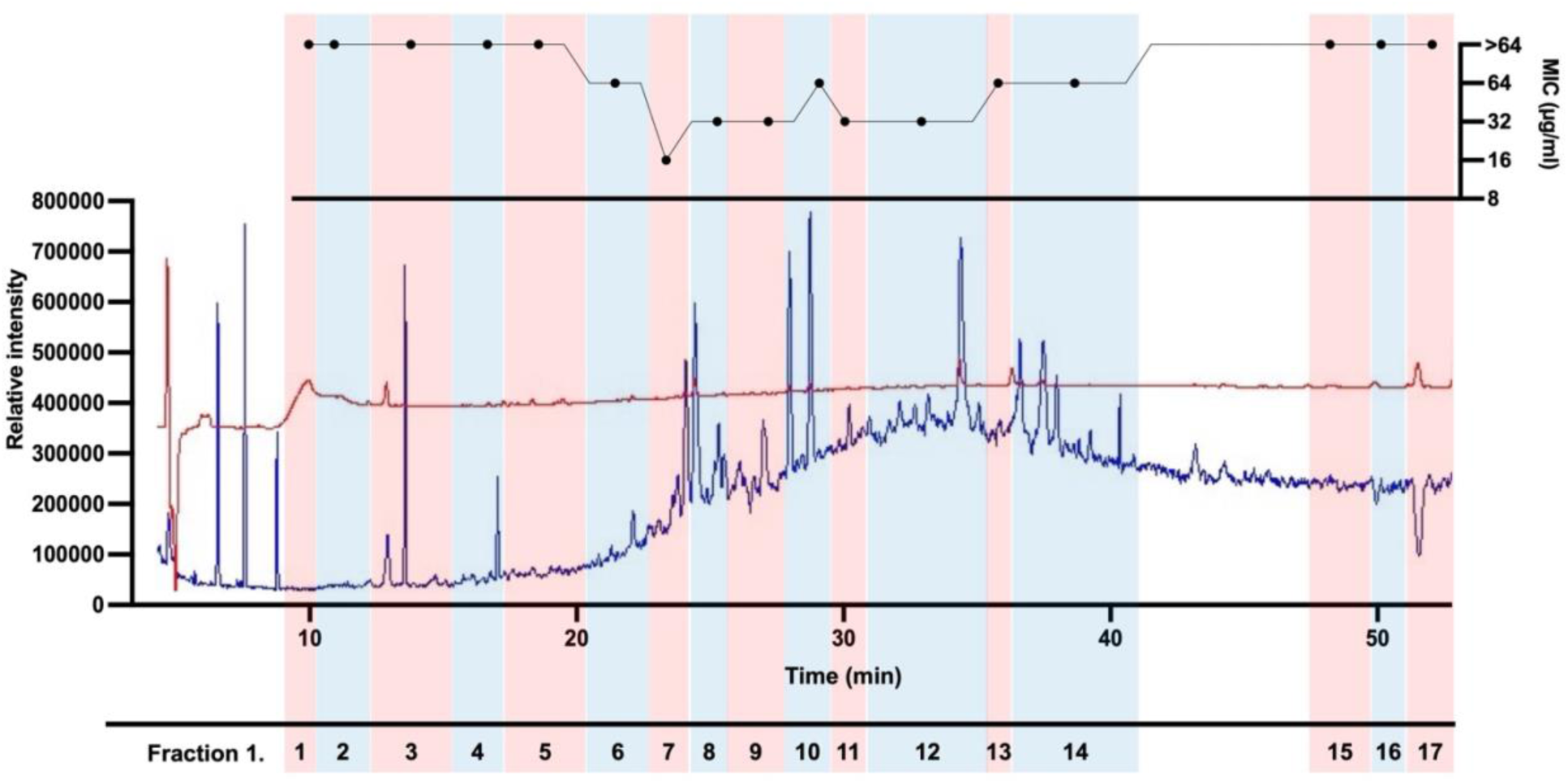
CE-MNAAK fraction 1.7 displays the lowest MIC against *E. coli ΔbamBΔtolC*. MNAAK crude extract was first fractionated using Prep-TLC, followed by further fractionation of Fraction 1 using HPLC with an ACN and H_2_O gradient. Fractions (1.1 to 1.17) were tested against *E. coli ΔbamBΔtolC* in an MIC assay at a maximal concentration of 64 µg/ml and 1.25% MeOH. Top panel represents the fraction’s MIC (mode of 3 biological replicates with 3 technical replicates each). Bottom panel represents LRMS (ESI+) Total Ion Chromatogram (TIC) (blue) and wavelength at 214 nm (red) of compounds in each fraction. Each fraction is highlighted in alternating colours, and numbers indicate the numbered subfractions generated from Fraction 1.

An initial search in the CHEMNet-base database (39) did not yield any compound with a m/z of 208. This led us to reanalyze the MNAAK genome sequence using the less stringent “loose” settings in antiSMASH and identify an 11-gene BGC with a high (0.84) similarity score (Figure 6A) to a putative polyketide biosynthesis cluster of bacterial tetrahydropyridine alkaloids. The biosynthetic products were recently described in *Pseudomonas koreensis* (40) and were termed koreenceines. Koreenceine A (HRMS (ESI+) *m/z* [C14H25N]+ calcd: 208.2065), is one of the four compounds characterized to be produced by this *kecA-K* gene cluster (40, 41). Homologues of the genes *kecA* to *kecJ* were present in the MNAAK genome (74.36% -92.57% amino acid sequence identity) but *kecK* was not (Figure 6A). The high homology to the koreenceine BGC and the detection of a putative bioactive compound with m/z compatible with koreenceine A suggested that the latter may be one of the antibacterial compounds in the MNAAK crude extract.

**Figure 6.**
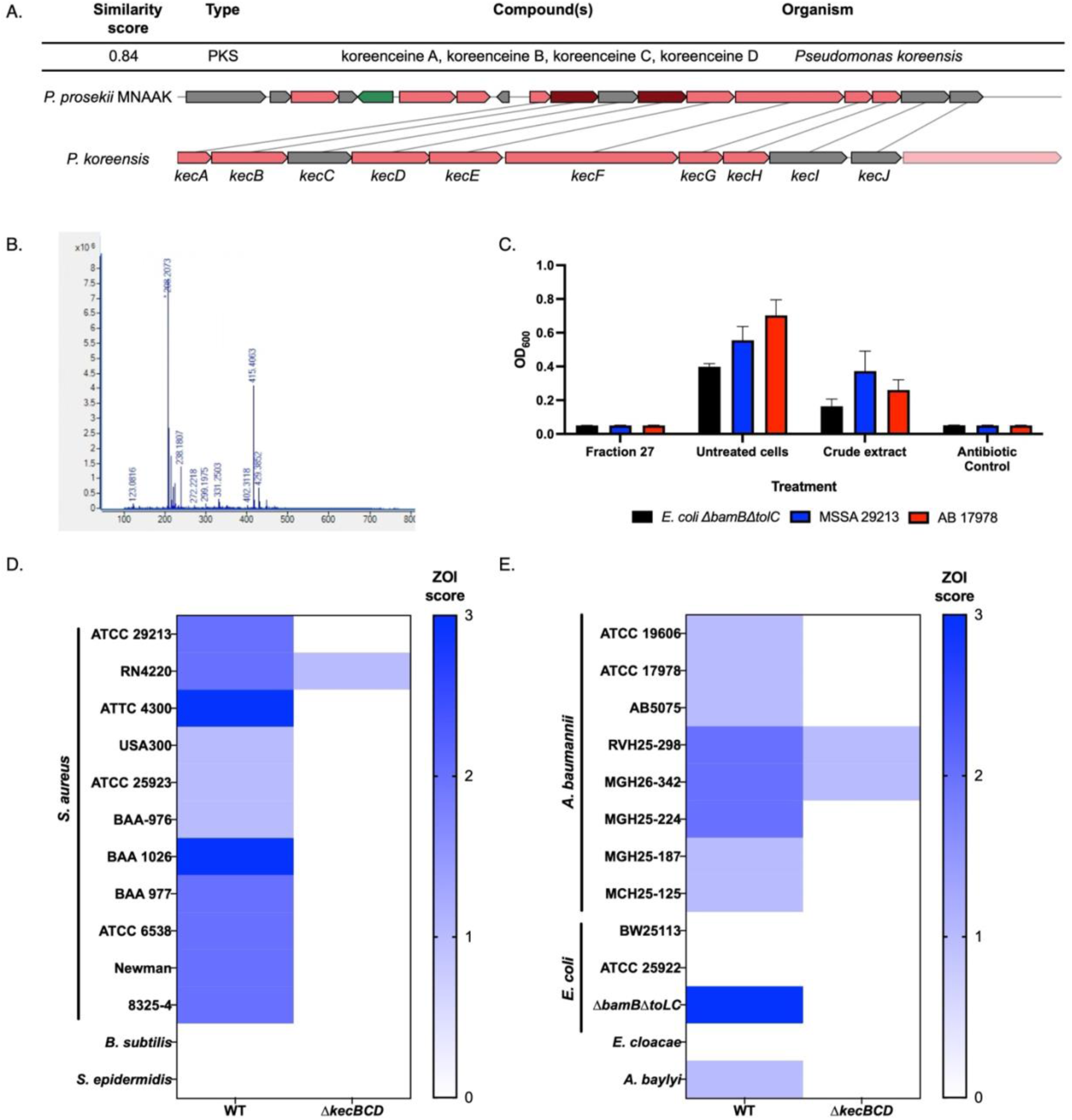
Identification of the putative biosynthetic gene cluster responsible for MNAAK antibacterial activity. **A.** AntiSMASH output of predicted BGC from MNAAK with a similarity score of 0.84 to the koreenceine BGC from *P. koreensis* containing genes *kecA – kecJ* (Burgundy: core biosynthetic genes, pink: additional biosynthetic genes, green: regulatory genes, grey: other genes). **B.** MNAAK crude extract was fractionated using RPLC on an HP C18 column with H_2_O and ACN. HRMS (ESI+) TIC of peak from fraction 27 putatively containing koreenceine A, (calcd: 208.2065; found: 208.2073). **C**. Fraction 27 was dried and resuspended in DMSO and tested against *E. coli* Δ*bamB* Δ*tolC*, *S. aureus* ATCC 29213, and *A. baumannii* ATCC 17978. Results are shown as mean (+/-SEM) from N=3 biological replicates. **D.** Antibacterial activity measured as ZOI in co-culture stamping assay of WT and Δ*kecBCD* MNAAK against **D.** Gram-positive and **E.** Gram-negative bacteria. Results in D and E are shown as the mode of N=6 from 3≤ independent experiments.

To confirm the chemical identity of the m/z 208 antibacterial compound(s) in CE_MNAAK, we sought to purify the compounds by Reversed Phase Liquid Chromatography (RPLC). After one round of fractionation using flash column chromatography, we retained 55 fractions for testing of antibacterial activity. Fraction 27 (HRMS (ESI+) *m/z* [C14H25N]+ calcd: 208.2065; found: 208.2073) (Figure 6B) displayed inhibitory activity against *E. coli* Δ*bamB* Δ*tolC*, *S. aureus* ATCC 29213, and *A. baumannii* ATCC 17978 (Figure 6C). Unfortunately, attempts to further purify koreenceine A from this fraction or from CE_MNAAK using flash column chromatography, HPLC, normal phase chromatography, and LH20 size exclusion chromatography were unsuccessful due to difficulties with compound separation and degradation.

### Construction of a koreenceine knockout mutant

To confirm whether the bioactive compounds in CE_MNAAK were produced by the putative koreenceine BGC, we constructed a koreenceine deletion mutant in the *P. prosekii* MNAAK strain to knockout the *kecB*, *kecC*, and *kecD* genes, and asked whether the loss of the koreenceine biosynthesis would abrogate its antibacterial activity. The *kecB* and *kecD* genes encode for putative ß-ketoacyl synthases required for the synthesis of koreenceine, while *kecC* encodes for a partial β-ketoacyl synthase with a conserved thiolase domain (40). As control, we confirmed the loss of the putative koreenceine A in the crude extract generated from Δ*kecBCD* strain by low resolution MALDI-TOF MS (Figure S5). MNAAK and its isogenic Δ*kecBCD* mutant were tested in co-culture against a panel of 26 bacterial tester strains, including 11 *S. aureus* and 8 *A. baumannii* human pathogenic strains. Compared to WT MNAAK, the Δ*kecBCD* mutant displayed reduced antibacterial activity against all *S. aureus* and *A. baumannii* strains, with a notable complete loss of activity against 10 *S. aureus* strains, 6 *A. baumannii* strains, *E. coli* Δ*bamB*Δ*tolC,* and *A. baylyi.* (Figure 6D & E). Neither the WT nor the Δ*kecBCD* mutant strains had antibacterial activity against other Gram-positive bacteria, namely *B. subtilis* and *S. epidermidis*, or against the Gram-negative *E. cloacae* and wild-type *E. coli* strains.

## Discussion

In this study, we developed an antibacterial discovery pipeline to screen, prioritize, and characterize the antibacterial activity of a collection of CHA bacterial isolates. This pipeline involved screening of soil samples from various CHA environments, integrating the Tiny Earth studentsourcing activities for the initial screen for antibacterial activity, followed by dereplication, whole genome analysis, and chemical characterization of bioactive crude extracts. From this CHA bacterial isolate collection we identified 11 Arctic isolates that produced potentially novel antibacterial compounds, prioritized 5 crude extracts of interest, and determined the putative BGC responsible for activity in one extract. The dominant genus in our collection was *Pseudomonas* (44.60%). While the prevalence of this genus in our collection appeared high, other studies of Arctic sediment and soil samples also reported this genus as most abundant.(42–45). However, these studies did not pre-screen for antibacterial activity and used a mix of culture-based and culture-independent techniques to obtain genomic DNA, which may account for the differences observed between these studies and ours. We note that approximately one-third (106 of 352) of the isolates in our collection come from the Gypsum Hill hummock soil sample. The high representation of isolates from this sampling site is likely due to the fact that this location was the only sampling site for the 2021 student cohort. Excluding the 2021 cohort, samples from the freshwater lake sediment and biofilm, and the enriched gas bubble pond sediment resulted in the largest number of CHA isolates that demonstrated antibacterial activity (26 isolates each), potentially suggesting that these environments are enriched in antimicrobial-producing bacteria. However, since the total number of isolates from each sampling location were not recorded, we cannot conclude whether the ratio between isolates with and without antibacterial activity differed between these sampling sites.

We incorporated the Tiny Earth studentsourcing science initiative into this study by incorporating students from an Introduction to Microbiology undergraduate lab course to isolate and screen bacteria from Arctic environmental samples. Tiny Earth, like other successful discovery-based research courses (46–49), aims to inspire students to address the AMR crisis while alleviating the labour-intensive process of bacterial isolation and screening (20). With efforts ongoing, over 43 institutions have submitted samples to the Tiny Earth program resulting in over 6,000 isolates with antimicrobial activity that have been publicly identified (21), and early reports are emerging of antimicrobial compounds such as labradorin 1 with anti-nematodal activity (50). Other citizen science antibiotic discovery initiatives have focused on expanding the environmental sampling efforts, but the bacterial isolation and screening remained in the hands of researchers (51–55). For example, the Swab and Send program reported hundreds of strains with activity against several human pathogens (51). While these initiatives have significant pedagogical value and raise awareness among students and the general public, their ability to discover new antibacterial compounds remains to be demonstrated, with only one novel compound discovered through the program to our knowledge (56).

Following initial isolation and screening by the Tiny Earth students, 167 isolates were assessed by the ARP for dereplication of which only 18 passed. The high attrition rate (89%) suggests that many Arctic bacteria are producing compounds similar to known antibiotics in clinical use. Whether this is due to an inherent ability to produce these conventional antibiotics, or influenced by other factors, for example, acquired from bacteria introduced from other environments, remains to be determined.

From our pipeline, the 11 unique Arctic isolates prioritized following screening and dereplication belong to three different genera (*Pseudomonas, Serratia*, and *Bacillus*), with *Pseudomonas* as the dominant genus. All three genera represented among the prioritized Arctic isolates are known to produce antibacterial compounds such as phenazines and mupirocin (*Pseudomonas*) (57–59), carbapenem and bacteriocins (*Serratia*) (60–63), and macrolactin, and benzoids (*Bacillus*) (64).

We recognize that our culturing methods introduced a culture bias for isolates that are thermotolerant and fast-growing on rich medium compared to culture-based isolation at low temperatures (4 - 18°C) for longer incubation time, and on minimal media, or culture-independent analyses (42, 43, 65). In this study, all samples were cultured on TSA media and incubated for one week at room temperature, and 8 out of 9 isolates identified from cryo-iPlate samples belonged to the *Pseudomona*s genus. In contrast, our group’s previous work culturing similar Arctic samples with cryo-iPlate cultured on minimal media (R2A) and incubated for 2 weeks for isolation following a 3-month *ex situ* incubation at 5°C (23) identified genera not identified in our current study. Our culturing methods were thus adapted for the studentsourcing workflow but likely led us to overlooking slow growing isolates, which have been shown to be promising reservoirs of novel antibacterial natural products (66, 67).

While long incubation may not be readily compatible with teaching labs, our screening pipeline could be improved upon for future iterations. Incorporating the *E. coli* Δ*bamB* Δ*tolC* strain into the initial screening could ensure that all isolates that move to the dereplication step can be screened using the ARP, whose strains are in the *E. coli ΔbamB ΔtolC* background (22). Additionally, a more targeted preliminary screening step using bioreporters (68, 69) for antibacterial mode of action can help gain additional insight into the inhibitory activity of isolates and help prioritize isolates of interest.

Gram-negative bacteria possess a highly impermeable outer membrane (OM) that contains selective OM porins and a variety of efflux pumps that expel xenobiotics out of the cell. These properties thus present a considerable barrier to antibacterial compounds and can hinder screening efforts (70). In this study, we carried out several antibacterial assays using as positive control the hyperpermeable and efflux-deficient *E. coli* Δ*bamB* Δ*tolC* strain, a genetically engineered mutant rendered hypersusceptible to antibiotics (25). Although the lipoprotein BamB is not essential for cell viability, it plays an important role in OM protein assembly and its deficiency leads to greater OM permeability (71). Meanwhile, TolC serves as an essential subunit of several efflux pumps whose deletion results in lack of antibiotic export (72). With these mutations, *E. coli* becomes sensitive to several antibiotics that typically only have activity against Gram-positive bacteria, such as erythromycin (macrolide), clindamycin (lincosamide), and rifampicin (rifamycin) (22). Such hyperpermeable and efflux-deficient strains can thus be used to screen a wider range of compounds for antibacterial activity, including hydrophobic compounds that typically cannot pass through the Gram-negative OM (73). However, compounds identified through such an approach may not demonstrate activity against wild-type *E. coli*, as observed with our MNAAK strain (Figure 6E).

The CE_MNAAK crude extract was prioritized as it displayed the most potent antibacterial activity against several *S. aureus* strains. Following the identification of a putative BGC with high homology to the koreenceine BGC and the presence of a compound compatible with koreenceine A (m/z 208.2073, calcd 208.2065) in our active fractions, we sought to determine whether this compound contributed to the antibacterial activity of MNAAK. Although our efforts to purify koreenceine A were unsuccessful, similar difficulties with compound separation and degradation were observed by Lozano et al. (40).

Using a different approach to examine whether koreenceine A was responsible for the antibacterial activity of the MNAAK isolate, we generated a Δ*kecBCD* knockout mutant by deleting the three key biosynthesis genes of the putative koreenceine BGC, namely *kecB* and *kecD* genes that encode for a β-ketoacyl synthase (KS*a*) and *kecC* gene that encodes for a partial β-ketoacyl synthase with a conserved thiolase domain (KSβ). (40). KS*a* and KSβ form a heterodimer complex with KS*a* required to catalyze the decarboxylative Claisen condensations between the malonyl-acyl carrier protein and the thioester intermediate, and KSβ functions as the chain length determining factor (74). It therefore follows that without KS*a* from *kecB* and *kecD*, synthesis of koreenceine would be disrupted, as suggested by the loss of the putative koreenceine A MALDI-TOF signal in the Δ*kecBCD* crude extract. This model is also indirectly supported by previous work by Lozano et al. which showed that a transposon mutant inserted within *kecD* resulted in a loss of antibacterial activity against *F. johnsoniae* (40).

We confirmed that the Δ*kecBCD* mutant lost activity against nearly all *S. aureus* (n=10 of 11) and *A. baumannii* (n=6 of 8) strains, but did however note that it had diminished but residual activity against methicillin-susceptible *S. aureus* RN4220 strain and two *A. baumannii* strains. The laboratory strain *S. aureus* RN4220 contains multiple mutations that affect its fitness, growth, and other cellular processes likely to alter its response to antibiotics (75). The two *A. baumannii* strains are clinical isolates with resistance to cefazolin. Together, these findings suggest that the *kecABCDEFGHIJ* BGC synthesizes compounds responsible for dominant antibacterial activity observed with MNAAK, but additional bioactive compound(s) are likely produced. Koreenceine A is likely the bioactive compound synthesized by MNAAK *kecABCDEFGHIJ* BGC but was challenging to purify, as previously reported by Lozano et al (40). Testing of a synthesized version of the compound would further confirm its antibacterial spectrum of activity and mechanism of action. Further bioactivity-guided fractionation of the Δ*kecBCD* mutant crude extract would also allow identification of other antibacterial compound(s) produced by MNAAK.

The mechanism of action of koreenceine A is unknown. A previous study has shown that in polymicrobial community, *P. koreensis*-produced koreenceine modulates gene expression in the translation, post-translational modification, protein turnover, and chaperones categories in both *F. johnsoniae* and *B. cereus* (76). Previously, antibacterial activity of koreenceine had only been demonstrated against *F. johnsoniae* and several other bacteriodetes from the soybean rhizosphere, with koreenceine A as the most likely major inhibitory compound (40). Although the koreenceine BGC has been observed within the *Pseudomonas* phylogeny (40), it is surprising that a compound produced by a *Pseudomonas* from Wisconsin, USA and associated with activity against rhizosphere bacteria is also produced by *Pseudomonas* from arid active layer permafrost soil. This suggests that koreenceine A must be beneficial and active in completely different habitats if it is conserved in *Pseudomonas* bacteria in the CHA environment.

In this study, we describe an expanded spectrum of koreenceine A to multiple strains of two bacterial human pathogens *S. aureus* and *A. baumannni*. The additional activity against both Gram-negative and Gram-positive pathogens suggests that this compound may have previously unknown broad spectrum activity. Koreenceine A is a chemical analog of γ-coniceine, a toxic plant alkaloid produced by poison hemlock (78, 79) that shares a common core aromatic ring. While this raises concern about potential non-specific toxic effects, it is interesting that koreenceine A had no observed antibacterial activity against several Gram-positive (*B. subtilis* and *S. epidermidis*) and Gram-negative (*K. pneumoniae, E. cloacae*, and WT *E. coli*) bacterial species tested.

Several recent studies have analyzed bacterial genomes from various soil environments and Arctic regions such as the Arctic Ocean and Glacier-fed streams (80–83). With an average of 14 BGCs per genome (Table 1), the Arctic isolates in our study contained a greater number of BGCs per genome compared to the average (2 – 8 BGCs per genome) observed in these studies. However, we note that our study only sequenced Arctic isolates with demonstrated antibacterial activity that passed dereplication, thus representing a selected set of isolates. Our analysis also used a more recent versions of antiSMASH (v8.0.1) (29), which contains a larger database of BGCs compared to older versions (v5.1.2 and v6.1.1) used in some of the previously published studies (80, 83).

The two most prevalent classes of BGCs identified among our prioritized Arctic isolates are non-ribosomal peptide synthetases (NRPS) and ribosomally synthesised and post-translationally modified peptide products (RiPP) (Figure S3). NRPSs have the ability to incorporate a vast range of substrates and synthesize highly diverse compounds, many of which are pharmaceutically relevant and include antibiotics such as vancomycin, polymyxin B, and bacitracin (84, 85). While RiPPs have not yet yielded the same pharmaceutical success as NRPSs, several antibacterial RiPPs have already been discovered (66, 86–92). Additionally, their large diversity of post-translational modifications result in enhanced compound stability and bioactivity which makes them attractive as potential sources of novel antibiotics (93). Varying abundance levels of BGCs encoding for NRPSs and RiPPs have been observed in polar bacteria (80, 94–96). Interestingly, no NRPS sequences were recovered from metagenomic analyses from samples collected ∼300 km east of our sampling site on Axel-Heiberg Island, possibly indicating that the prevalence of such BGC classes may be specific to sample site or habitats (94).

## Conclusion

In this study, we developed a pipeline for the isolation, screening, and characterization of bacteria from environmental samples and created a collection of CHA bacterial isolates with antibacterial activity. By incorporating the Tiny Earth studentsourcing initiative into our workflow, we provided valuable educational experiences and successfully identified bacterial strains with antibacterial activity. Our work suggests that many Arctic isolates produce compounds similar to known antibiotics, though others produce lesser known, or potentially novel antibacterial compounds. The continued expansion and screening of the collection will help provide more insight into the antibacterial compounds, both known and unknown, that are produced by Arctic bacteria.

## Author Contributions

EF, SG, and DN designed the study and experiments, and interpreted the data. EF, ML, GP, LB, DS carried out experiments. EF, MN, KK and FLM performed fractionation on CE_MNAAK. PL performed sequencing of the WT MNAAK isolate under the supervision of JS. KA and GDW contributed to the study design, methods, data analysis and supervision. LW contributed arctic samples. SG supervised the Tiny Earth teaching lab. EF and DN wrote the main manuscript. All authors reviewed and approved the manuscript.

## Acknowledgments

We would like to acknowledge the students of McGill’s MIMM212 course for performing the initial isolation and screening of Arctic bacteria. We would like to acknowledge McGill AMR Centre Microbial Library (McGill University, Montréal, Canada, https://bit.ly/ML_access;) for providing the clinical isolates obtained this study. We would also like to acknowledge Anne-Laure Larocque and the Proteomics and Molecular Analysis Platform for their support and access to the mass spectrometry. This research was supported by funding from the Natural Sciences and Engineering Research Council (RGPIN-2019-06408 to DN), the Canada Graduate, Fonds de Recherche Quebec Nature Technologie, and the Canadian Network for Antimicrobial Resistance (CAN-AMR-Net) Trainee scholarships (to EF), and the McGill Interdisciplinary in Initiative Infection and Immunity (to KA and DN).

We would especially like to acknowledge our colleague and co-author Dr. Lyle Whyte, whose research in the Canadian High Arctic advanced our knowledge of microbes in harsh environments and astrobiology. His dedication to his research and mentorship inspired us all and he will be deeply missed.

## Data availability

Raw sequencing reads have been deposited in the Sequence Read Archive (SRA) under the BioProject ID PRJNA1526223. All data generated for this study are available upon request to the corresponding author, Dao Nguyen

## Notes

https://www.ncbi.nlm.nih.gov/sra/PRJNA1526223

### Competing Interest Statement

The authors have declared no competing interest.

