## Supplemental file for "Exploring the antibiotic-producing capabilities of Canadian High Arctic bacteria"

**Supplemental Methods**

**Whole genome sequencing (WGS) and analysis**

Genomic DNA was purified using the DNeasy Blood & Tissue Kit (Qiagen) according to the manufacturer's protocol for all isolates. Hybrid WGS for 2018 Arctic isolates were performed by the SeqCenter (Pittsburgh, PA). Paired-end (2x 150 bp) short-read sequencing was done on the Illumina NovaSeq 6000 platform following standard Illumina library preparation and PCR amplification. Raw sequences were demultiplexed, quality-filtered, and adapter-trimmed using bcl-convert (v4.1.5). Long-read sequencing was done using the Oxford Nanopore Technologies (ONT) platform. Libraries were prepared without PCR amplification, using the Native Barcoding Kit (ONT, SQK-NBD114.24) in combination with the NEBNext Companion Module (E7180L) following the manufacturer's protocols. Sequencing was conducted on MinION Mk1B or GridION (ONT) instruments equipped with R10.4.1 flow cells. Guppy (v6.4.6) was used for super accuracy base calling (SUP model), demultiplexing, and adapter trimming (dna_r10.4.1_e8.2_400bps_modbases_5mc_cg_sup.cfg). Sequencing reads were trimmed using Trimmomatic (v0.38.1) (1) (Illumina reads) and Porechop (0.2.4+galaxy0) (2) (Nanopore reads). Hybrid assembly was performed using tools available in the open source, web-based platform Galaxy (https://usegalaxy.org/) with de novo assembly using Unicycler (v 0.5.0+galaxy1) (3) under default settings.

For long-read sequencing of WT MNAAK, genomic DNA was cleaned up using PuroMAG™ NGSprep Magnetic Beads (Luna Nanotech) and quantified using a Qubit fluorometer. 200 ng gDNA was sequenced using the Nanopore Rapid Barcoding 96 v14 (ONT) on a R10.4.1M PromethION flow cell. The Dorado base caller integrated into MinKNOW v 23.07.5 (Oxford Nanopore Technologies) was used for base calling and demultiplexing raw sequencing to FASTQ files (dna_r10.4.1_e8.2_400bps_sup@v4.3.0). WGS analysis was processed using tools available in the open source, web-based platform Galaxy (https://usegalaxy.org/). Sequencing reads were trimmed using Porechop (0.2.4+galaxy0) (2), de novo genome assembly was performed using Flye (v 2.9.1+galaxy0) (4).

For long-read sequencing of 2019 and 2021 isolates, as well as the MNAAK Δ*kecBCD* mutant, sequencing was performed using ONT platform by Plasmidsaurus. Sequencing was conducted on PromethION or MinION (ONT) instruments equipped with R10.4.1 flow cells. Dorado v4.3 Super-Accurate Basecalling with default Q10 quality filtering was used for base calling (SUP model), and demultiplexing raw sequencing to FASTQ files (dna_r10.4.1_e8.2_400bps_sup@v4.3.0). De novo assembly was performed by Plasmidsaurus with quality filtering using Filtlong (v0.2.1) (5), and assembly using Autocycler(6) using Flye (v2.9.6+)(4), HiFiasm (7), and Plassembler (v1.8.0+) (8).

Whole genome sequences of Arctic isolates were compared for average nucleotide identity (ANI) using FastANI (9). Isolates with ANI ≥ 99.99% were considered identical strains and only one of the isolates was retained for further characterization. All isolates with ANI < 99.99% were considered different strains and continued in the pipeline (Table S4). Taxonomic classification was done using the Microbial Genomes Atlas (MiGA) (10).

**<u>MALDI-TOF analysis of WT and Δ*kecBCD* extracts</u>**

WT and Δ*kecBCD* MNAAK were grown under optimized conditions in 500 mL of R2A (Teknova) media in a 2 L flask at 15˚C for 24 h and 250 RPM. Following the established extraction protocol, CE_MNAAK and CE_Δ*kecBCD*_MNAAK were resuspended to an approximate concentration of 1mg / mL in DMSO before 1 μL was spotted on the MALDI-TOF plate with 1 μL of 10 mg/ mL α-cyano-4-hydroxycinnamic acid reconstituted in 80% ACN. Fraction 27 was used as a positive control. Spectra were analyzed using the Bruker UltrafleXtreme in positive reflector mode with 5000 laser shots.

**Supplemental Tables**

**Table S1.Pathogen-relative tester species use in this study.**

| | | | |
|---|---|---|---|
| *Enterococcus raffinosus* ATCC 49464 | Tiny Earth isolate screening, pathogen relative of *Enterococcus faecium* | ATCC | (11) |
| *Bacillus subtilis* | Tiny Earth isolate screening | Handelsman Lab, University of Wisconsin-Madison | (11) |
| *Staphylococcus epidermidis* ATCC 14990 | Tiny Earth isolate screening, pathogen relative of *Staphylococcus aureus* | ATCC | (11) |
| *Escherichia coli* ATCC 11775 | Tiny Earth isolate screening, pathogen relative of *Klebsiella* species | ATCC | (11) |
| *Acinetobacter baylyi* ATCC 33305 | Tiny Earth isolate screening, pathogen relative of *Acinetobacter baumannii* | ATCC | (11) |
| *Pseudomonas putida* | Tiny Earth isolate screening, pathogen relative of *Pseudomonas aeruginosa* | Handelsman Lab, University of Wisconsin-Madison | (11) |
| *Enterobacter aerogenes* ATCC 51697 | Tiny Earth isolate screening, pathogen relative of *Enterobacter* species | Handelsman Lab, University of Wisconsin-Madison | (11) |

**Table S2. Taxonomic classification of Arctic isolates with antibacterial activity against pathogen-relative tester species.** Taxonomic classification of n=352 Arctic isolates by 16S rRNA gene sequencing. The species of the isolates was identified using nucleotide BLAST, with a threshold of minimum 87% match to known species.

| Genus | Number of Arctic isolates in each genus |
|---|---|
| *Pseudomonas* | 157 |
| Unknown | 36 |
| *Streptomyces* | 33 |
| *Bacillus* | 15 |
| *Flavobacterium* | 12 |
| *Janthinobacterium* | 7 |
| *Arthrobacter* | 6 |
| *Citrobacter* | 5 |
| *Paenibacillus* | 5 |
| *Pedobacter* | 4 |
| *Acinetobacter* | 3 |
| *Sphingomonas* | 3 |
| *Staphylococcus* | 3 |
| *Aeromonas* | 2 |
| *Chryseobacterium* | 2 |
| *Clostridium* | 2 |
| *Iodobacter* | 2 |
| *Klebsiella* | 2 |
| *Kocuria* | 2 |
| *Limosilactobacillus* | 2 |
| *Microbacteriaceae* | 2 |
| *Moraxella* | 2 |
| *Paenarthrobacter* | 2 |
| *Psychrobacter* | 2 |
| *Salinibacterium* | 2 |
| *Shewanella* | 2 |
| *Acidovorax* | 1 |
| *actinomycetes* | 1 |
| *Agreia* | 1 |
| *Bosea* | 1 |
| *Corynebacterium* | 1 |
| *Enterobacter* | 1 |
| *Enterococcus* | 1 |
| *Kitasatospora* | 1 |
| *Leifsonia* | 1 |
| *Leucobacter* | 1 |
| *Lysinibacillus* | 1 |
| *Lysobacter* | 1 |
| *Marinobacter* | 1 |
| *Micrococcus* | 1 |
| *Morganella* | 1 |
| *Mucilaginibacter* | 1 |
| *Nocardia* | 1 |

| | |
|---|---|
| *Pectobacterium* | 1 |
| *Planococcus* | 1 |
| *Priestia* | 1 |
| *Pseudoarthrobacter* | 1 |
| *Rheinheimera* | 1 |
| *Rhodococcus* | 1 |
| *Rugamonas* | 1 |
| *Serratia* | 1 |
| *Stenotrophomonas* | 1 |
| *Yersinia* | 1 |

**Table S3. Antibiotic Resistance Platform (ARP) strains used in this study for dereplication.**

| | | | |
|---|---|---|---|
| *Escherichia coli* ΔbamBΔtolC BW25113 pGDP3 aph(3")-Ia | ARP strain, resistant to aminoglycosides (streptomycin) | Wright Lab, McMaster University | (12) |
| *Escherichia coli* ΔbamBΔtolC BW25113 pGDP3 rmtB | ARP strain, resistant to aminoglycosides (2- deoxystreptamine) | Wright Lab, McMaster University | (12) |
| *Escherichia coli* ΔbamBΔtolC BW25113 pGDP3 apmA | ARP strain, resistant to aminoglycosides (apramycin) | Wright Lab, McMaster University | (12) |
| *Escherichia coli* ΔbamBΔtolC BW25113 pGDP3 aph(9)-Ia | ARP strain, resistant to aminoglycosides (spectinomycin) | Wright Lab, McMaster University | (12) |
| *Escherichia coli* ΔbamBΔtolC BW25113 pGDP1 NDM-1 | ARP strain, resistant to β-lactams (penicillin, cephalosporin, carbapenam) | Wright Lab, McMaster University | (12) |
| *Escherichia coli* ΔbamBΔtolC BW25113 pGDP4 ermC | ARP strain, resistant to lincosamides, macrolides, type B streptogramins | Wright Lab, McMaster University | (12) |
| *Escherichia coli* ΔbamBΔtolC BW25113 pGDP3 vatD | ARP strain, resistant to type A Streptogramins | Wright Lab, McMaster University | (12) |
| *Escherichia coli* ΔbamBΔtolC BW25113 pGDP1 STAT | ARP strain, resistant to streptothricin | Wright Lab, McMaster University | (12) |
| *Escherichia coli* ΔbamBΔtolC BW25113 pGDP4 tet(A) | ARP strain, resistant to tetracyclines (tetracycline) | Wright Lab, McMaster University | (12) |
| *Escherichia coli* ΔbamBΔtolC BW25113 pGDP3 CAT | ARP strain, resistant to chloramphenicols (chloramphenicol) | Wright Lab, McMaster University | (12) |
| *Escherichia coli* ΔbamBΔtolC BW25113 pGDP1 fosA | ARP strain, resistant to fosfomycins | Wright Lab, McMaster University | (12) |
| *Escherichia coli* ΔbamBΔtolC BW25113 pGDP3 arr | ARP strain, resistant to rifamycins | Wright Lab, McMaster University | (12) |

| | | | |
|---|---|---|---|
| *Escherichia coli* Wild Type BW25113 pGDP1 MCR-1 | ARP strain, resistant to polymyxins | Wright Lab, McMaster University | (12) |
| *Escherichia coli* ΔbamBΔtolC BW25113 pGDP1 uvrA | ARP strain, resistant to echinomycins | Wright Lab, McMaster University | (12) |
| *Escherichia coli* ΔbamBΔtolC BW25113 pGDP1 vph | ARP strain, resistant to tuberactinomycins (viomycin) | Wright Lab, McMaster University | (12) |
| *Escherichia coli* ΔbamBΔtolC BW25113 fhuB mutant | ARP strain, resistant to sideromycins (albomycin) | Wright Lab, McMaster University | (12) |
| *Escherichia coli* ΔbamBΔtolC BW25113 pGDP1 qnrA1 | ARP strain, resistant to fluoroquinolones (levofloxacin) | Wright Lab, McMaster University | (12) |

**Table S4. Average nucleotide identity of Arctic isolates that passed the ARP.** Average nucleotide identity (ANI) of genome sequences of Arctic isolates were compared using FastANI to identify identical strains. Strains with an ANI > 99.99% were considered identical and subsequently only one of each identical group was used in the next steps of the study. Identical pairs highlighted in red.

| | 107_2018 | 109_2018 | 26_2018 | 49_2018 | 57_2018 | 58_2018 | 5_2018 | 113_2021 | 2_2019 | 4_2019 | 46_2021 | 67_2021 | 74_2021 | 80_2021 | 110_2021 | MNAAK | 8_2021 |
|---|---|---|---|---|---|---|---|---|---|---|---|---|---|---|---|---|---|
| 109_2018 | 99.998 | | | | | | | | | | | | | | | | |
| 26_2018 | 99.999 | 99.996 | | | | | | | | | | | | | | | |
| 49_2018 | 100.000 | 99.998 | 99.999 | | | | | | | | | | | | | | |
| 57_2018 | 100.000 | 99.998 | 99.999 | 100.000 | | | | | | | | | | | | | |
| 58_2018 | 100.000 | 99.998 | 99.999 | 100.000 | 100.000 | | | | | | | | | | | | |
| 5_2018 | 90.001 | 89.955 | 89.966 | 89.945 | 90.003 | 90.005 | | | | | | | | | | | |
| 113_2021 | 81.988 | 82.020 | 81.981 | 81.992 | 81.987 | 81.987 | 81.844 | | | | | | | | | | |
| 2_2019 | 86.097 | 86.141 | 86.127 | 86.135 | 86.099 | 86.101 | 86.066 | 82.466 | | | | | | | | | |
| 4_2019 | 85.710 | 85.707 | 85.676 | 85.702 | 85.710 | 85.707 | 85.461 | 81.802 | 88.458 | | | | | | | | |
| 46_2021 | < 80 | < 80 | < 80 | < 80 | < 80 | < 80 | < 80 | < 80 | < 80 | < 80 | | | | | | | |

| | | | | | | | | | | | | | | | | | |
|---|---|---|---|---|---|---|---|---|---|---|---|---|---|---|---|---|---|
| 67_2021 | < 80 | < 80 | < 80 | < 80 | < 80 | < 80 | < 80 | < 80 | < 80 | < 80 | < 80 | | | | | | |
| 74_2021 | < 80 | < 80 | < 80 | < 80 | < 80 | < 80 | < 80 | < 80 | < 80 | < 80 | 95.316 | < 80 | | | | | |
| 80_2021 | < 80 | < 80 | < 80 | < 80 | < 80 | < 80 | < 80 | < 80 | < 80 | < 80 | 95.327 | < 80 | 99.998 | | | | |
| 110_2021 | < 80 | < 80 | < 80 | < 80 | < 80 | < 80 | < 80 | < 80 | < 80 | < 80 | < 80 | < 80 | < 80 | < 80 | | | |
| MNAAK | 87.306 | 87.297 | 87.303 | 87.285 | 87.303 | 87.306 | 87.049 | 81.640 | 85.974 | 85.630 | < 80 | < 80 | < 80 | < 80 | < 80 | | |
| 8_2021 | < 80 | < 80 | < 80 | < 80 | < 80 | < 80 | < 80 | < 80 | < 80 | < 80 | 99.287 | < 80 | 95.266 | 95.324 | < 80 | < 80 | |
| 49_2021 | < 80 | < 80 | < 80 | < 80 | < 80 | < 80 | < 80 | < 80 | < 80 | < 80 | 100.000 | < 80 | 95.290 | 95.305 | < 80 | < 80 | 99.255 |

### Table S5. Human pathogen screening strains used in this study.

| | | | |
|---|---|---|---|
| *Escherichia coli ΔbamBΔtolC* BW25113 | hyperpermeable, efflux-deficient *E. coli* | Wright Lab, McMaster University | (13) |
| *Staphylococcus aureus* RN4220 | Human pathogen tester strain | Heinrichs Lab, Western University | (14) |
| *Staphylococcus aureus* ATCC 29213 | Human pathogen tester strain | ATCC | (15) |
| *Staphylococcus aureus* ATCC 43300 | Human pathogen tester strain | ATCC | |
| *Staphylococcus aureus* USA300 | Human pathogen tester strain | Yau Lab, University of Toronto | (16) |
| *Acinetobacter baumannii* ATTC 19606 | Human pathogen tester strain | ATCC | (17) |
| *Acinetobacter baumannii* AB5075 | Human pathogen tester strain | Ramage Lab, University of Washington | (18) |
| *Klebsiella pneumoniae* ATCC 700603 | Human pathogen tester strain | ATCC | (19) |
| *Klebsiella pneumoniae* C0859 | Human pathogen tester strain | Wright Lab, McMaster University | |
| *Pseudomonas aeruginosa* ATCC 27853 | Human pathogen tester strain | ATCC | (20) |
| *Pseudomonas aeruginosa* DNC231 | Human pathogen tester strain, "*P. aeruginosa* AMT0023-30" | Burns Lab, University of Washington | (21) |
| *Staphylococcus aureus* ATCC 25923 | Human pathogen tester strain | ATCC | (22) |
| *Staphylococcus aureus* BAA 976 | Human pathogen tester strain | Yansouni Lab, McGill University | |
| *Staphylococcus aureus* BAA 1026 | Human pathogen tester strain | Yansouni Lab, McGill University | (23) |
| *Staphylococcus aureus* BAA 977 | Human pathogen tester strain | Yansouni Lab, McGill University | (24) |

| | | | |
|---|---|---|---|
| *Staphylococcus aureus* ATCC 6538 | Human pathogen tester strain | ATCC | (25) |
| *Staphylococcus aureus* Newman | Human pathogen tester strain | Heinrichs Lab, Western University | (26) |
| *Staphylococcus aureus* NCTC 8325-4 | Human pathogen tester strain | Heinrichs Lab, Western University | (27) |
| *Acinetobacter baumannii* ATCC 17978 | Human pathogen tester strain | ATCC | (28) |
| *Acinetobacter baumannii* RVH25-298 | Clinical isolate | McGill AMR Centre Microbial Library | (29) |
| *Acinetobacter baumannii* MGH26-342 | Clinical isolate | McGill AMR Centre Microbial Library | (29) |
| *Acinetobacter baumannii* MGH25-224 | Clinical isolate | McGill AMR Centre Microbial Library | (29) |
| *Acinetobacter baumannii* MGH25-187 | Clinical isolate | McGill AMR Centre Microbial Library | (29) |
| *Acinetobacter baumannii* MCH25-125 | Clinical isolate | McGill AMR Centre Microbial Library | (29) |
| *Enterobacter cloacae* C0794 | Clinical isolate | Wright Lab, McMaster University | |
| *Enterobacter cloacae* V3082440 | Clinical isolate | Longtin Lab, McGill University | |
| *Enterobacter cloacae* MGH24-011 | Clinical isolate | McGill AMR Centre Microbial Library | (29) |
| *Enterobacter cloacae* RVH24-043 | Clinical isolate | McGill AMR Centre Microbial Library | (29) |
| *Enterobacter cloacae* ATCC 13047 | Human pathogen tester strain | ATCC | (30) |
| *Escherichia coli* WT BW25113 | Human pathogen tester strain | Wright Lab, McMaster University | (31) |
| *Escherichia coli* ATCC 25922 | Human pathogen tester strain | ATCC | (32) |
| *Streptococcus pyogenes* MCH24-006 | Clinical isolate | McGill AMR Centre Microbial Library | (29) |
| *Streptococcus anginosus* MGH26-352 | Clinical isolate | McGill AMR Centre Microbial Library | (29) |
| *Streptococcus pyogenes* MGH25-068 | Clinical isolate | McGill AMR Centre Microbial Library | (29) |
| *Streptococcus anginosus* MGH24-044 | Clinical isolate | McGill AMR Centre Microbial Library | (29) |

**Table S7. Prioritized Arctic isolates in this study**

| Strain | Relevant Information | Source location | Reference |
|---|---|---|---|
| *Pseudomonas mandelii* 57-2018 | Isolated from gravel soil | This study | |
| *Pseusdomonas prosekii* MNAAK | Isolated from arid active layer permafrost soil | Whyte Lab, McGill University | (33) |
| *Pseudomonas wadenswilerensis* 113-2021 | Isolated from Gypsum Hill hummock Gypsum Hill hummock active layer permafrost soil | This study | |
| *Pseudomonas koreensis* 2-2019 | Isolated from Tupirvik Beach Sediment | This study | |
| *Bacillus pumilus* 110-2021 | Isolated from Gypsum Hill hummock active layer permafrost soil | This study | |
| *Bacillus subtilis* 67-2021 | Isolated from Gypsum Hill hummock active layer permafrost soil | This study | |
| *Serratia marcescens* 8_2021 | Isolated from Gypsum Hill hummock active layer permafrost soil | This study | |
| *Pseudomonas rossensis* 5-2018 | Isolated from Arctic Willow rhizobium | This study | |
| *Serratia marcescens* 74-2021 | Isolated from Gypsum Hill hummock active layer permafrost soil | This study | |
| *Pseudomonas baetica* 4-2019 | Isolated from ice wedge polygon permafrost (10 in. Depth) | This study | |
| *Serratia marcescens* 49-2021 | Isolated from Gypsum Hill hummock active layer permafrost soil | This study | |

### Supplemental Figures

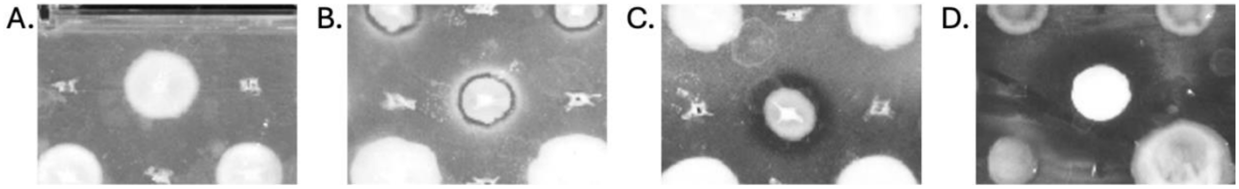


**Figure S1. Different ZOI sizes producing by Arctic bacteria during co-culture stamping assay.** Antibacterial activity of Arctic isolates was measured in a co-culture stamping assay by ranking the ZOI 0 (none), 1 (small), 2 (medium), and 3 (large). Representative images of different ZOI **A.** ZOI = 0 (none). **B.** ZOI = 1 (small). **C.** ZOI = 2 (medium). **D.** ZOI = 3 (large).

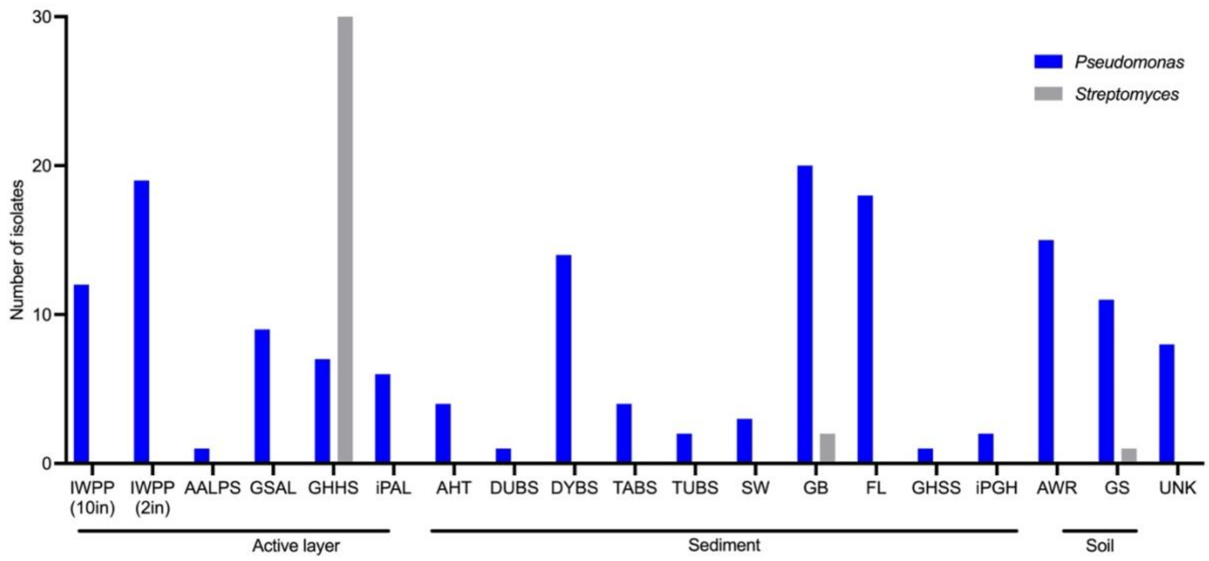


**Figure S2. Sampling location of Canadian High Arctic isolates**. Locations include ice wedge polygon active-layer permafrost at 10 inch depth (IWPP 10in), ice wedge active-layer polygon permafrost at 2 inch depth (IWPP 2in), arid active layer permafrost soil (AALPS), gravel soil active layer (GSAL), Gypsum Hill hummock active layer permafrost soil (GHHS), supplemented media active layer permafrost soil (iPAL), Assistance Bay Beach sediment (high tide) (AHT), Dump Beach sediment (DUBS), Dynamite Beach sediment (DYBS), Tank Beach sediment (TABS), Tupirvik Beach sediment (TUBS), Arctic Ocean (salt water) sediment (SW), enriched gas bubble pond sediment (GB), freshwater lake sediment and biofilm (FL), Gypsum Hill spring sediments (GHS), Gypsum Hill Spring outflow channel sediment (GHSS), supplemented media Gypsum Hill Spring cryo-iplate plugs (iPGH), Arctic Willow (*Salix arctica*) rhizobium (AWR), and gravel soil (GS). Isolates with unknown location were coded as UNK.

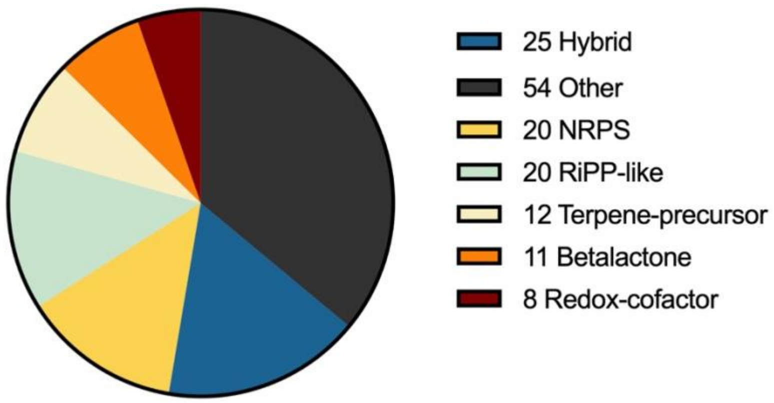

**Figure S3. Breakdown of the top 5 BGC classes from 11 prioritized Arctic isolates.** Genomes of 11 prioritized Arctic isolates were analyzed using antiSMASH v8.0.4 using the "relaxed" setting, resulting in 151 identified BGCs. The top 5 BGC classes: Non-ribosomal peptide synthetase (NRPS), ribosomally synthesized post-translationally modified-like (RiPP-like), terpene-precursor, beta-lactone, and redox-cofactor are shown. Hybrid BGCs were labelled as "hybrid" and BGC classes that were not part of the top five classes were labelled as "other."

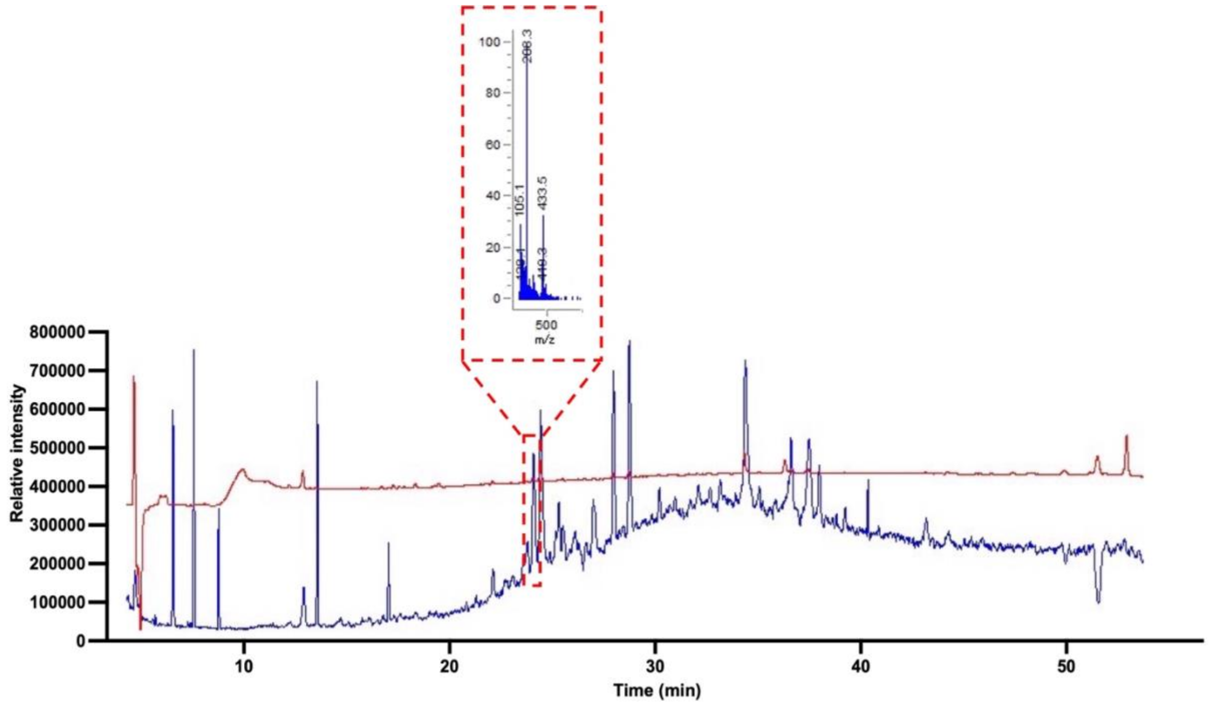


**Figure S4. Fraction 1.7 contains a compound with m/z of 208.3.** Total Ion Chromatogram (TIC, blue tracing) and at wavelength 214 nm (red tracing) using LR-MS of Fraction 1 generated from Prep-TLC. Inset shows corresponding fraction 1.7 (most active fraction) TIC peak indicating a m/z of 208.3.

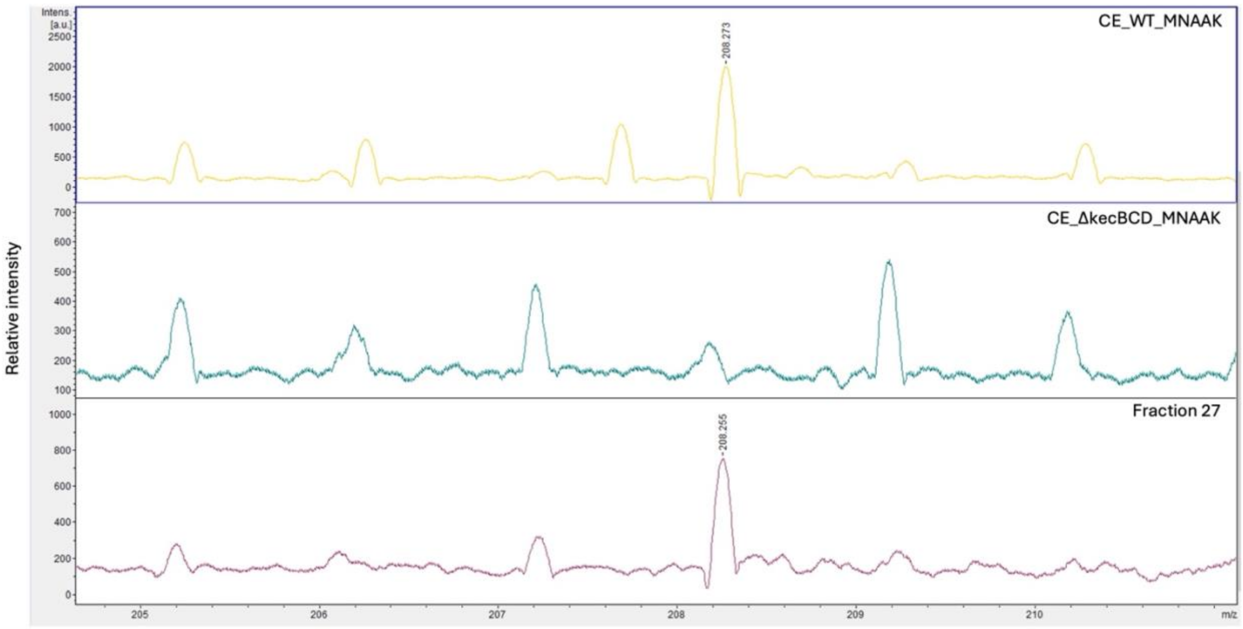


**Figure S5. Koreenceine A is not present in the Δ*kecBCD* crude extract.** Ethyl acetate crude extracts from WT and Δ*kecBCD* bacterial supernatant were analyzed by Low-Resolution Matrix-Assisted Laser Desorption/Ionization Time-of-Flight Mass Spectrometry (MALDI-TOF). Fraction 27 was used as a control.
